# The astrocyte α1-adrenoreceptor is a key component of the neuromodulatory system in mouse visual cortex

**DOI:** 10.1101/2023.05.02.538934

**Authors:** Jérôme Wahis, Cansu Akkaya, Andre M. Kirunda, Aline Mak, Karen Zeise, Jens Verhaert, Hayk Gasparyan, Sargis Hovhannisyan, Matthew G. Holt

## Abstract

Noradrenaline (norepinephrine) is known to modulate many physiological functions and behaviors. In this study, we tested to what extent astrocytes, a type of glial cell, participate in noradrenergic signaling in mouse primary visual cortex (V1). Astrocytes are essential partners of neurons in the central nervous system. They are central to brain homeostasis, but also dynamically regulate neuronal activity, notably by relaying and regulating neuromodulator signaling. Indeed, astrocytes express receptors for multiple neuromodulators, including noradrenaline, but the extent to which astrocytes are involved in noradrenergic signaling remains unclear. To test whether astrocytes are involved in noradrenergic neuromodulation in mice, we knocked down the major noradrenaline receptor in astrocytes, the α1A- adrenoreceptor. Using this model, we found that the α1A-adrenoreceptor is involved in triggering intracellular calcium transients in astrocytes, which are generally thought to underlie astrocyte function. To test if impaired α1A-adrenoreceptor signaling in astrocytes affected the function of neuronal circuits in V1, we used electrophysiological measurements and found that noradrenergic signaling through astrocyte α1A-adrenoreceptor controls the basal level of inhibition and regulates plasticity in V1 by potentiating synaptic responses in circuits involved in visual information processing.

**MAIN POINTS:**

- The α1A-adrenoreceptor (α1A-NAR) is the major α1-NAR in primary visual cortex (V1)
- α1A-NAR signaling triggers astrocyte Ca^2+^ responses in V1.
- Astrocyte α1A-NARs play a key role in regulating basal inhibitory transmission and are crucial for LTP induction in V1

## INTRODUCTION

Astrocytes are a major cell type in the mammalian central nervous system (CNS), with functions ranging from homeostatic regulation to synaptic modulation (Verkhratsky and Nedergaard, 2018). Astrocytes are generally considered electrically non-excitable cells. However, changes in their local environment, principally neuronal activity and the resulting release of neurotransmitters and/or neuromodulators, elicits transient fluctuations in their intracellular calcium ([Ca^2+^]). These [Ca^2+^] transients are thought to regulate astrocyte functions, including the release of gliotransmitters, which reciprocally regulate the activity of neural circuits (Rusakov, 2015; Goenaga et al., 2023). These key functions have been confirmed in numerous studies, using a variety of animal models, underlying the importance of astrocytes for correct neural circuit function and the generation of specific behaviors (Nagai et al., 2021).

Astrocytes integrate neuronal activity through a wide repertoire of receptors, including those for a number of neuromodulators (Verkhratsky and Nedergaard, 2018). One neuromodulator particularly studied in relation to its action on astrocytes is noradrenaline (NA, also known as norepinephrine). NA afferences in neocortex project from the main noradrenergic nucleus - the locus coeruleus (LC) (Schwarz and Luo, 2015; Poe et al., 2020). The firing patterns of LC neurons are thought to predict changes in behavioral states, with phasic burst firing being associated with responses to salient stimuli (Berridge and Waterhouse, 2003). Interestingly, noradrenergic projections form few synapses, with release of NA typically occurring from axonal varicosities directly opposed to astrocytic processes (Vizi et al., 2010). Based on this anatomical evidence, NA has been proposed to signal principally through volume transmission, with astrocytes as major targets (Hirase et al., 2014; Wahis and Holt, 2021).

Cortical astrocyte [Ca^2+^] responses to phasic NA release have been reported, with one adrenoreceptor (NAR) appearing central to these responses - α1-NAR (Bekar et al., 2008; Ding et al., 2013; Paukert et al., 2014; Slezak et al., 2019; Oe et al., 2020; Reitman et al., 2023). To date, however, it is not clear whether this is a direct effect of NA on astrocytes, an indirect effect resulting from NA acting on neurons or a combination of both. This knowledge gap stems, at least in part, from the complexity of the α1-NAR family. This is comprised of three distinct G_q_-coupled G-protein-coupled receptors: α1A-, α1B- and α1D- NAR, each of which is transcribed from an independent gene (Adra1a, Adra1b and Adra1d, respectively). This has greatly complicated the use of conventional gene ablation approaches to investigate α1-NAR function in astrocytes. However, recent reports using astrocyte-specific conditional knock-out (cKO) of Adra1a have provided compelling evidence that expression of α1A-NAR in astrocytes is essential for NA effects in the CNS. Kohro and colleagues found that astroglial α1A-NAR is necessary for noradrenergic modulation of mechanical pain in spinal cord (Kohro et al., 2020), while the Paukert lab confirmed a central role of α1A-NAR in generating [Ca^2+^] transients in (cerebellar) astrocytes (Ye et al., 2020; Salinas-Birt et al., 2023). Crucially, Reitman and colleagues demonstrated that α1A-NAR-mediated astrocyte responses act as an important negative feedback mechanism, limiting the effects of NA on cortical circuit activity (Reitman et al., 2023). Indeed, while increased arousal, and the associated phasic NA release, initially desynchronizes cortical networks, consistent with the known effects of NA (Polack et al., 2013; McGinley et al., 2015), the authors observed that α1A-NAR signaling in astrocytes was important to counteract effects of arousal on neuronal activity, restoring circuits to a more synchronized state, notably after the effects of phasic NA release on neuronal activity peaked (Reitman et al., 2023). With this study, we sought to test whether α1A-NAR signaling is important to generate [Ca^2+^] transients in cortical astrocytes and aimed to better understand how it modulates neuronal network activity on different timescales.

Previous studies looking at the role of NA in cortex reported transient increases in astrocyte [Ca^2+^] in supragranular layers (layers I-III) (Bekar et al., 2008; Ding et al., 2013; Paukert et al., 2014; Slezak et al., 2019; Oe et al., 2020; Reitman et al., 2023), consistent with the pattern of Adra1a expression observed by Reitman and colleagues (Reitman et al., 2023). Therefore, we chose to focus on the same layers in mouse primary visual cortex (V1), as astrocytes in this brain region have been shown to exert powerful effects on the plasticity of neuronal circuits involved in visual information processing (Müller and Best, 1989; Chen et al., 2012; Pankratov and Lalo, 2015; Monai et al., 2016; Hennes et al., 2020; Ribot et al., 2021). Using our previously published single cell RNA-sequencing (scRNA-seq) astrocyte database, we confirmed that Adra1a is the most highly expressed (α1-)NAR transcript in cortical astrocytes (Figure 1), and designed a panel of shRNAs allowing astrocyte-specific gene knockdown (KD) (Fellmann et al., 2013). Using a combination of optical imaging, patch-clamp and multi-electrode array recordings in brain slices, we found that α1A-NAR activation in V1 astrocytes is important in the generation of [Ca^2+^] responses in astrocytes and noradrenergic neuromodulation of cortical circuits. Our results indicate that astrocytes are integral components of neural circuits in V1, acting as key relays of neuromodulator activity to control circuit function.

**Figure 1.**
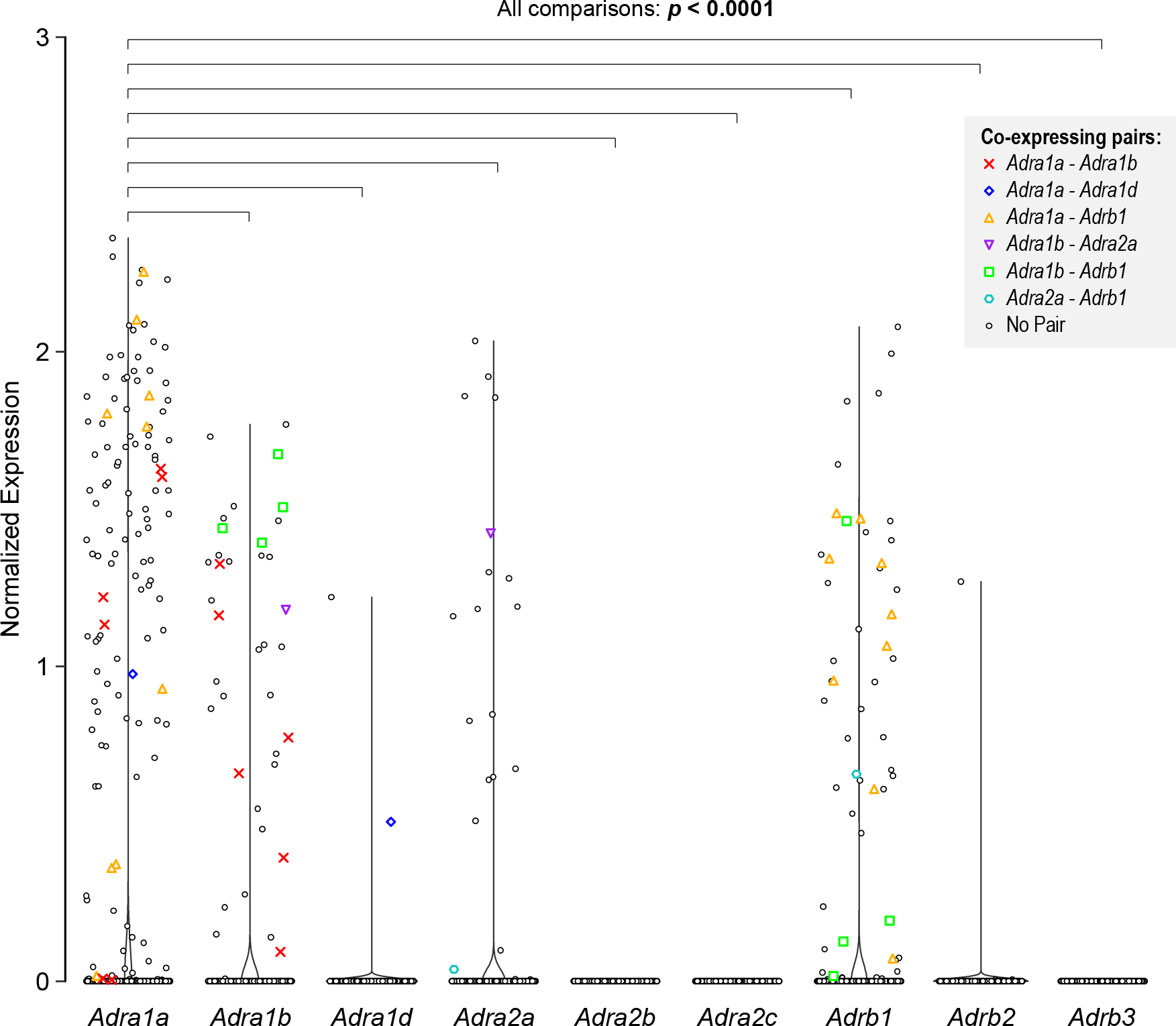
α1A-NAR is the most prevalent subtype of NAR in mouse cortex. Violin plots representing the normalized mRNA expression in cortical astrocytes of the 9 NAR genes, based on previously published single cell data (Batiuk et al., 2020). Adra1a, Adra1b, Adra1d, Adra2a, Adra2b, Adra2c, Adrb1, Adrb2 and Adrb3 encode for the α1A-, α1B-, α1D-, α2A-, α2B-, α2C-, β1-, β2- and β3-NAR, respectively. Among the 990 cortical astrocytes contained in the dataset, 128 (12.93%) were positive for Adra1a, 38 (3.84%) for Adra1b, 2 (0.20%) for Adra1d, 20 (2.02%) for Adra2a, 0 for Adra2b and Adra2c, 54 (5.45%) for Adrb1, 3 (0.30%) for Adrb2 and 0 for Adrb3. Each symbol in each gene group represents a single astrocyte and all of the 990 astrocytes analyzed are displayed in each gene group. Colored symbols represent astrocytes which co-express a defined pair of NARs, as specified in the figure key. Black circles represent astrocytes expressing only one or no NARs. Statistics displayed above the graph indicate p-values obtained from Dunn’s tests adjusted with the Bonferroni method. These tests were performed following a significant Kruskal-Wallis test. Statistical test parameters are summarized in Table S1. The number of astrocytes co-expressing defined NAR pairs, as well as the values of Spearman correlation tests and their corresponding p-values, are given in Table S2.

## MATERIAL AND METHODS

### Animal experiments

All experiments were approved by the Ethical Research Committee of the KU Leuven and were in accordance with the European Communities Council Directive of 22 September 2010 (2010/63/EU) and with the relevant Belgian legislation (KB of 29 May 2013). Fgfr3-iCreER^T2^ mice (Young et al., 2010) were obtained from William D. Richardson (University College London) under a Material Transfer Agreement. Both male and female adult mice, aged between P71–P262, were used. Mice were housed under a 12 hours light / 12 hours dark cycle, with access to food and water ad libitum.

### AAV-mediated, astrocyte-specific shRNA expression strategy

We based our gene knockdown strategy on the pAV-CreOn-shRNA (FLEx) plasmids from Vigene Biosciences (Schnütgen et al., 2003), packaged in AAV with a PHP.B capsid, as this vector system was previously found to transduce astrocytes efficiently, following stereotaxic injection (Rincon et al., 2018), with little to no induction of cell reactivity (Chatterjee et al., 2021). Following transduction, in the absence of Cre recombinase, the pAV-CreOn-shRNA vector expresses the fluorescent marker mStrawberry in all transduced cells, due to incorporation of the strong ubiquitous cytomegalovirus (CMV) promoter in the vector expression cassette. However, in cells expressing active Cre recombinase, inversion of the CMV promoter leads to co-expression of an EGFP marker and shRNA, allowing easy cell identification (Figure 2A). The system we used was based on a miR-30-based shRNA backbone, and all shRNA sequences were designed using Vigene Biosciences’s proprietary shRNA design software.

**Figure 2.**
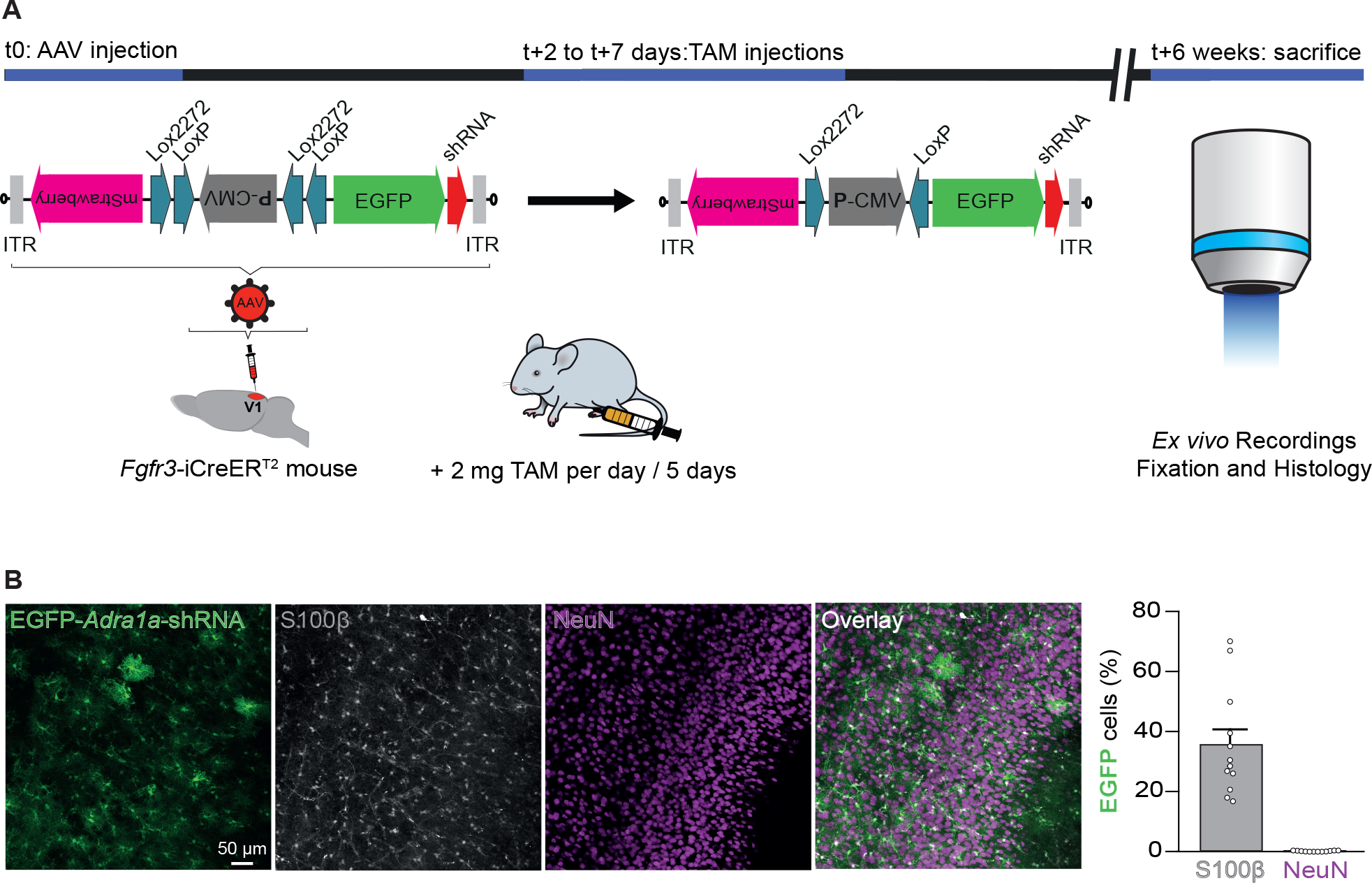
AAV-mediated EGFP-shRNA expression specifically in V1 astrocytes. A. Experimental scheme used to induce α1A-NAR knockdown in V1 astrocytes of adult mice. shRNAs were delivered into V1 via stereotaxic injection of an AAV-based vector into Fgfr3-iCreER^T2^ mice, which expressed a tamoxifen inducible Cre recombinase in (cortical) astrocytes. All AAVs used in this study were based on use of a FLEx expression cassette. Following initial vector injection, the orientation of the cytomegalovirus promoter (P-CMV) drives the expression of the fluorescent protein mStrawberry in all transduced cells. Following intraperitoneal tamoxifen injections, Cre-mediated recombination occurs between Lox sites, switching the orientation of the P-CMV, which then drives co-expression of EGFP and a miR-30-based short hairpin RNA (shRNA). B. (left) Following tamoxifen injections, the EGFP-shRNA construct (green) is expressed specifically in astrocytes in V1. S100β (grayscale): astrocytic marker; NeuN (magenta): neuronal marker. (Right) averaged proportions of astrocytes and neurons expressing EGFP- shRNA. n = 4 mice, 3 sections per mouse, white circles indicate individual section averages.

**Figure 3.**
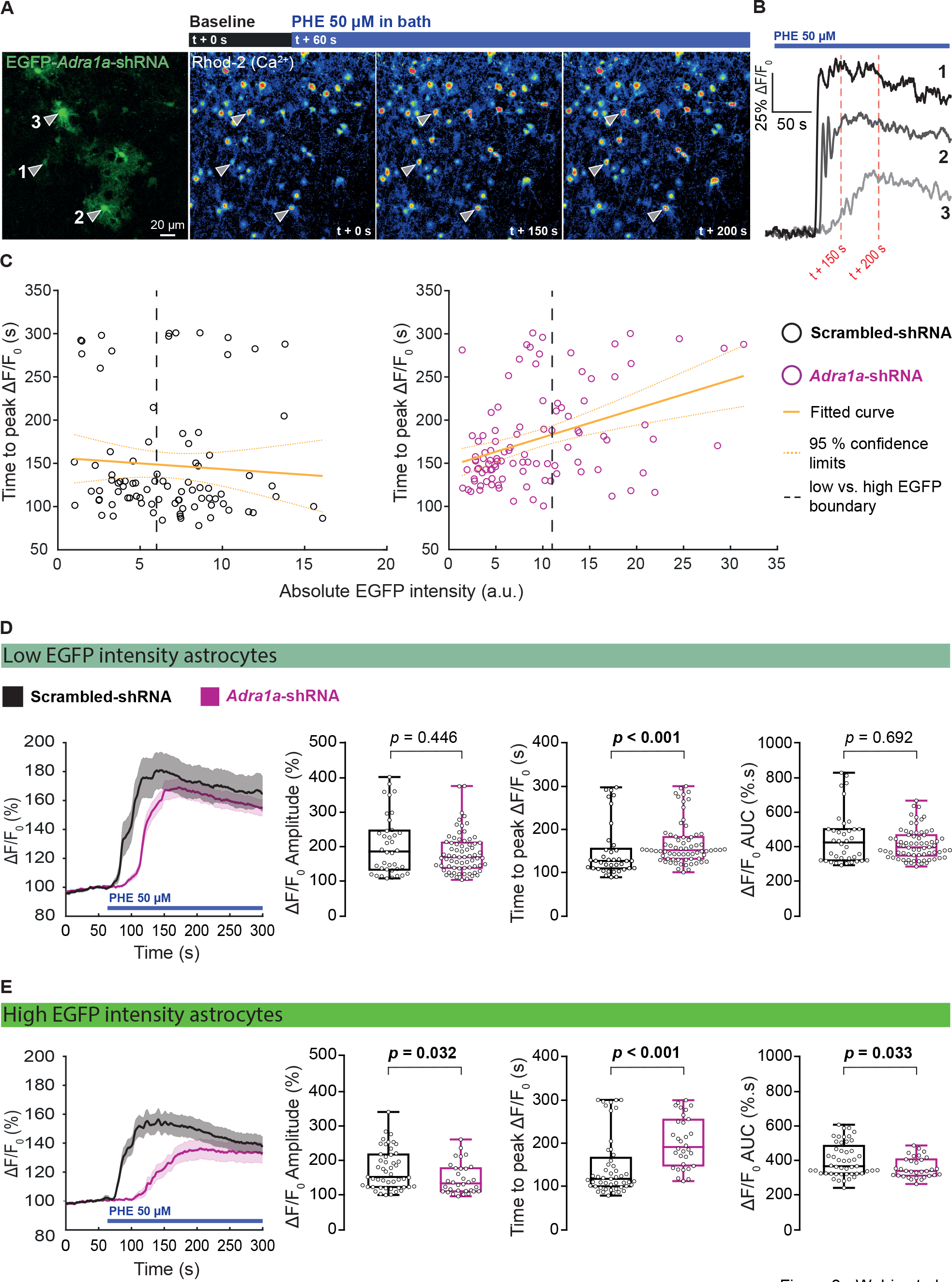
α1A-NAR signaling triggers astrocyte responses in V1. A. Calcium imaging of Rhod-2-loaded astrocytes expressing the Adra1a-shRNA construct. (Left) example picture of EGFP-Adra1a-shRNA positive astrocytes and (right) pseudo-color images of the Rhod-2 channel for the same cells at baseline and following phenylephrine (PHE) application at two different timepoints. The pseudo-color images represent an average of multiple frames acquired over a 10s period centered on the indicated timepoints. Arrowheads and numbers represent three cells of interest B. Example ΔF/F_0_ traces for cells of interest indicated in (A). The red dotted lines represent the timepoints around which the averaged pseudo-color images in (A) were obtained. C. Scatter plots of the time to peak for the PHE response as a function of the absolute EGFP fluorescence measured in shRNA expressing astrocytes (circles). The thick yellow line indicates a linear regression model fitted to the data; the dashed yellow lines indicate the 95% confidence intervals. The model parameters are indicated in the main text. The black dashed line indicates the chosen boundary segregating low and high EGFP intensity bins, used in the following figure panels. D-E (Left) averaged ΔF/F_0_ traces (solid line) and corresponding SEM (shaded area) for astrocytes expressing scrambled-shRNA (black) or Adra1a-shRNA (magenta). (Right) Properties of astrocyte responses to PHE. Cells were grouped as low or high EGFP intensity cells depending on their absolute EGFP fluorescence, as indicated in (C). ((D), scrambled-shRNA: n = 38 astrocytes; Adra1a-shRNA: n = 70 astrocytes; (E), scrambled-shRNA: n = 49 astrocytes; Adra1a-shRNA: n = 32 astrocytes). Scrambled-shRNA: n = 8 mice. Adra1a-shRNA: n = 7 mice. p-values for Wilcoxon rank sum test are indicated above the box-plot graphs and are displayed in bold when below 5%. White circles indicate values from individual astrocytes. All descriptive statistics and statistical test parameters are summarized in Table S1.

For maximum gene knockdown efficiency, we a generated three 3 separate AAVs each expressing a different Cre-dependent shRNA targeting an independent, non-overlapping section of the Adra1a mRNA.

The shRNA sequences were as follows: Adra1a-shRNA1:

5’-AGCGCGCCAGCACAGGTGAACATTTTAGTGAAGCCACAGATGTAAAATGTTCACCTGTGCTGGCG-3’

Adra1a-shRNA2:

5’-AGCGTGCCCTTCTCTGCCATCTTTGTAGTGAAGCCACAGATGTACAAAGATGGCAGAGAAGGGCA-3’

Adra1a-shRNA3:

5’-AGCGGATGAGACCATCTGCCAAATTAGTGAAGCCACAGATGTAATTTGGCAGATGGTCTCATCC-3’

A final injection mixture was created by mixing equal volumes of these AAVs. The final titer of each of the viral vectors in this mix was (in genome copies / ml): Adra1a-shRNA1: 6.30 x 10^12^; Adra1a-shRNA2: 5.20 x 10^12^, Adra1a-shRNA3: 1.17 x 10^13^.

As a control we generated a single AAV expressing a scrambled-shRNA sequence:

5’-AGCGCAACAAGATGAAGAGCACCAATAGTGAAGCCACAGATGTATTGGTGCTCTTCATCTTGTTG-3’

This vector had a final injected titer of 5.85 x 10^13^.

All experiment were conducted using a single production batch of each AAV, produced by Vigene Biosciences. When referring to vectors without distinguishing between the shRNAs expressed, we use the term “EGFP-shRNA” in the text. When referring to the AAV mixture targeting Adra1a, we use the terminology “Adra1a-shRNA”. Finally, when referring to the control vector, we use “scrambled-shRNA”.

To allow for astrocyte-specific expression, AAVs were injected into the Fgfr3-iCreER^T2^ mouse line (Young et al., 2010), in which astrocytes specifically express iCreER^T2^, allowing recombination at Lox sites following tamoxifen administration. Note that in our hands, some neurons expressed mStrawberry (data not shown), indicating they were transduced by the vector, but that promoter reorientation and EGFP expression were strictly dependent on Cre activity. Conversely, we never observed mStrawberry positive astrocytes 6 weeks following tamoxifen administration, indicating high recombination efficiency.

### AAV stereotaxic injections

Mice were anesthetized using isoflurane gas inhalation (4% in O_2_ for induction, 1.5% in O_2_ for maintenance). Mice were subsequently injected through a craniotomy into the primary visual cortex (V1) of both hemispheres. Coordinates used were 3.4 mm posterior to bregma, +/- 2.5 mm lateral to the midline, at a depth of 300 μm from the cortical surface. Viral vector solutions were injected via a polished glass capillary using a Nanoject II Auto-Nanoliter Injector (Drummond Scientific). A triple injection approach per craniotomy was chosen to reach an optimal volume of transduced area in V1, based on a maximal distance of 300 μm between injection sites along the A-P axis of the brain. Each injection had a total volume of 300 nl, which was injected over the course of 5 minutes. After each injection, the capillary was left in place for 90 s before being slowly retracted to minimize backflow of the AAV solution along the capillary tract, ensuring maximum transduction efficacy. Mice were administered analgesics during and after the procedure, following guidelines of the Ethical Research Committee of the KU Leuven.

### Tamoxifen administration

Tamoxifen (MP biomedicals) was dissolved in warm (50°C) corn oil to produce a stock solution of 10 mg/ml concentration. Intraperitoneal injections of 2 mg tamoxifen (per mouse) were performed every day for 5 consecutive days. Experiments started a minimum of six weeks following the first tamoxifen administration.

### Immunohistochemistry

To assess the cell-type specificity of our knockdown approach, tamoxifen treated mice were transcardially perfused with phosphate-buffered saline (PBS) followed by 4% paraformaldehyde solution in PBS. Brain regions containing the primary visual cortex were cut into 50 µm thick coronal sections using a VT100S vibratome (Leica Biosystems). Sampled sections were then blocked with 10% Normal Donkey Serum (NDS, Abcam) and 0.2% Triton X-100 (Sigma-Aldrich) in Tris-buffered saline (TBS) for 3h at room temperature. Next, sections were incubated overnight at 4°C in a TBS-based primary antibody solution, composed of: 10% NDS, 0.2% Triton X-100, mouse anti-S100β (1:1000 dilution, Sigma-Aldrich, S2532), rabbit anti-GFP (1:500 dilution, Synaptic Systems, 132002) or guinea pig anti-NeuN (1:400 dilution, Synaptic Systems, 266004). Following 6 x 10 min washes in TBS, sections were then incubated for 3 h at room temperature in a TBS-based secondary antibody solution, composed of 2.5% NDS, 0.2% Triton X-100, donkey anti-mouse Cy3 (1:200 dilution, Jackson ImmunoResearch, 715-165-150), donkey anti-rabbit Alexa Fluor 488 (1:200 dilution, Invitrogen, A21206) or donkey anti-guinea pig Cy5 (1:200 dilution, Jackson ImmunoResearch, 706-175-148). Following 6 x 10 min washes in TBS, sections were mounted onto Superfrost slides (ThermoFisher Scientific) and coverslipped in Fluoromount-G medium with DAPI (ThermoFisher Scientific). Stained sections were imaged with an SP8x confocal microscope (Leica Microsystems), equipped with standard excitation and emission filters. Images were acquired using a 20X objective (HC PL APO CS, 20X/0.70 NA), and exported as separate TIFF files. We imaged 25 to 30 consecutive confocal planes of 0.75 µm thickness in each section, in regions in close proximity to the injection site. We used the Fiji ‘Cell counter’ plugin (Schindelin et al., 2012) to manually count S100β, S100β + EGFP, NeuN and NeuN + EGFP positive cells across the acquired stacks, and made a ratio of the number of S100β + GFP / S100β and NeuN + GFP / NeuN cells counted. Identification of single cells was controlled for by the obligate presence of a single DAPI stained nucleus in each counted cell. We averaged the ratio obtained in each condition across all analyzed sections and expressed it as a percentage, with the result displayed in Figure 2B.

To label biocytin-filled neurons and EGFP-shRNA expressing astrocytes following patch-clamp recordings, the slices were swiftly transferred from the recording chamber to 500 µl of 4% paraformaldehyde solution in PBS, for 48 h at 4°C. The slices were then washed 3 x 5 min in PBS and stored at 4°C in PBS until use. For immunohistochemistry, the slices were incubated in blocking solution composed of 10% NDS and 0.5% Triton X-100 in PBS for 8 h at room temperature. Slices were then incubated for 65 h at 4°C in a PBS-based primary antibody and streptavidin containing solution, composed of 2.5% NDS, 0.5% Triton X-100, rabbit anti-GFP (1:500 dilution, Synaptic Systems, 132002) and Alexa Fluor 647-conjugated streptavidin, (1:200 dilution, Jackson ImmunoResearch, 016-600-084). Following 6 x 1 h washes in PBS, sections were incubated overnight at 4°C in secondary antibody solution, composed of 2.5% NDS, 0.2% Triton X-100 and donkey anti-rabbit Alexa Fluor 488 (1:200 dilution, Invitrogen, A21206). Following 6 x 1 h washes in PBS, sections were mounted onto Superfrost slides and coverslipped in Fluoromount-G medium with DAPI. Stained sections were imaged with an SP8x confocal microscope (Leica Microsystems) equipped with standard excitation and emission filters. Images were acquired using a 40X objective (HC PL APO CS2 40X/1.30 NA) and exported as separate TIFF files. A 20 μm thick Z-stack, consisting of 20 planes of 1 µm thickness centered around the patched neuron, was obtained - and its maximal projection used to generate the illustrative picture in Figure 4A. For the illustrative image displayed in Figure 5B, the same immunohistochemistry protocol was applied, but the Alexa Fluor 647 conjugated streptavidin was omitted and the image was acquired using a DM5500 microscope (Leica Microsystems), equipped with a 2.5X objective (N PLAN, 2.5X/0.07 NA). The image was exported to a TIFF file, and aligned with the brightfield image of the same slice obtained during multi-electrode array (MEA) recordings to produce the overlay picture displayed in Figure 5B.

**Figure 4.**
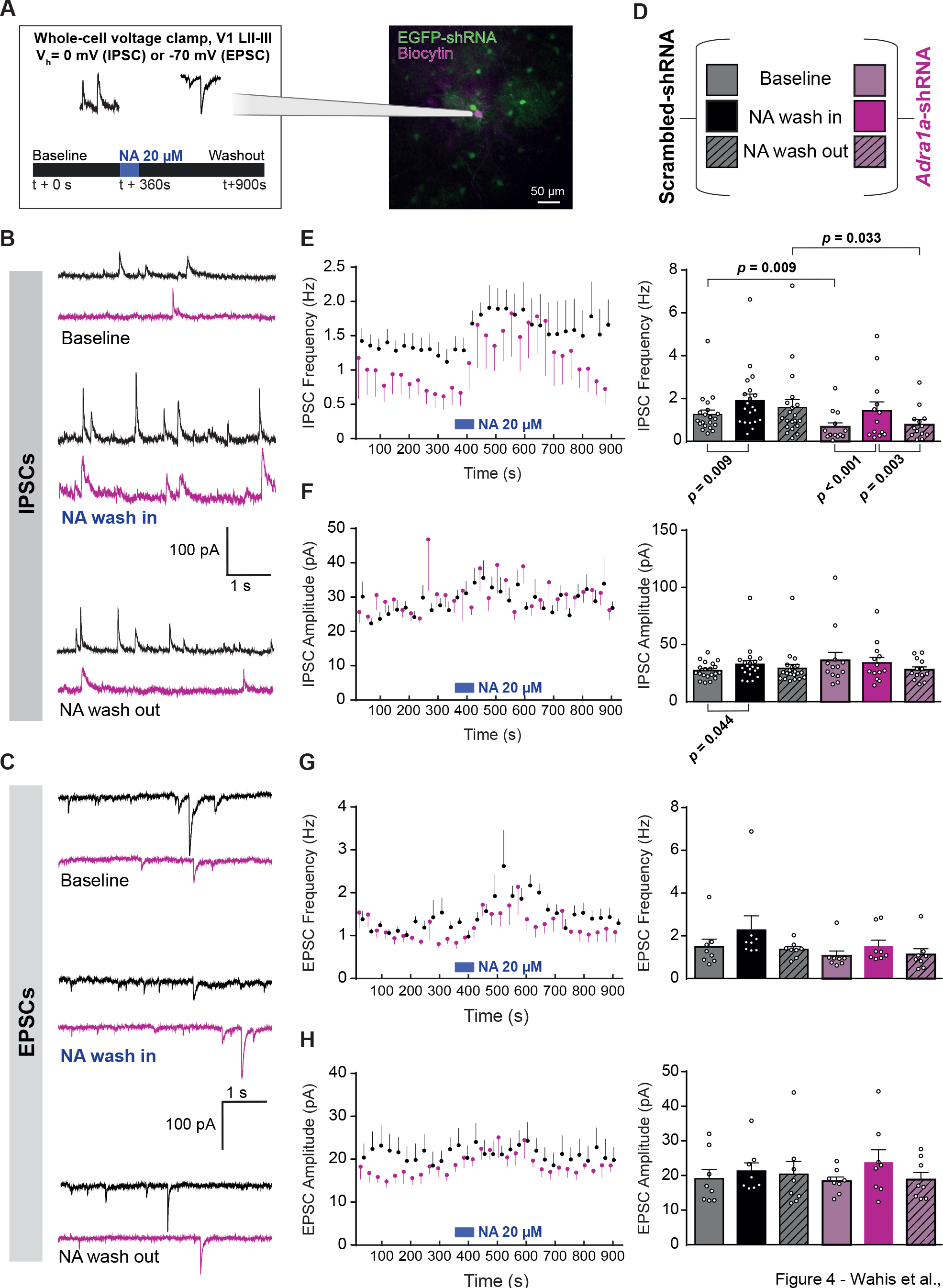
Astrocyte α1A-NAR plays a key role in regulating basal inhibitory transmission. A. (Left) Experimental scheme of the patch-clamp experiments presented in this figure. Neurons were patched in whole cell configuration and voltage-clamped at 0 or -70 mV, to isolate IPSCs or EPSCs, respectively. Following a 360 s baseline recording of spontaneous PSCs, NA was bath-applied to slices for 60 s and then left to wash out until the recording was stopped at t + 900 s. (Right) Neurons were patched in an area with visible EGFP-shRNA expression in V1 layer II-III astrocytes. In this example image, biocytin was added to the internal solution of the electrode to reveal the patched neuron, and its position embedded in a group of EGFP-shRNA expressing astrocytes. B. Example traces of IPSCs during baseline, NA wash in and NA wash out. C. Example traces of EPSCs during baseline, NA wash in and NA wash out. D. Legend key for the plots presented in consecutive panels. E. (Left) Time course of the change in IPSC frequency averaged in 30 s bins, vertical lines represent SEM. (Right) barplots of the IPSC frequency averaged over a 60 s time window for baseline (t = 240 – 300 s), NA wash in (t = 450 – 510 s) and NA wash out (t = 840 – 900 s) conditions. Error bars represent SEM. F. Same as (E), but for IPSC amplitude. G. Same as (E), but for EPSC frequency. H. Same as (E), but for EPSC amplitude. For IPSC measurements, scrambled-shRNA: n = 6 mice, 20 neurons; Adra1a-shRNA: n = 5 mice, 13 neurons. For EPSC measurements, scrambled-shRNA: n = 5 mice, 8 neurons; Adra1a-shRNA: n = 4 mice, 8 neurons. White circles indicate values from individual neurons. Statistics displayed above graphs indicate p-values for Wilcoxon rank sum test. Statistics displayed below graphs indicate p-values for Wilcoxon signed rank tests. Only p-values below 5% are displayed for figure clarity, p-values from other tests can be found in Table S1. All descriptive statistics and statistical test parameters are summarized in Table S1.

**Figure 5.**
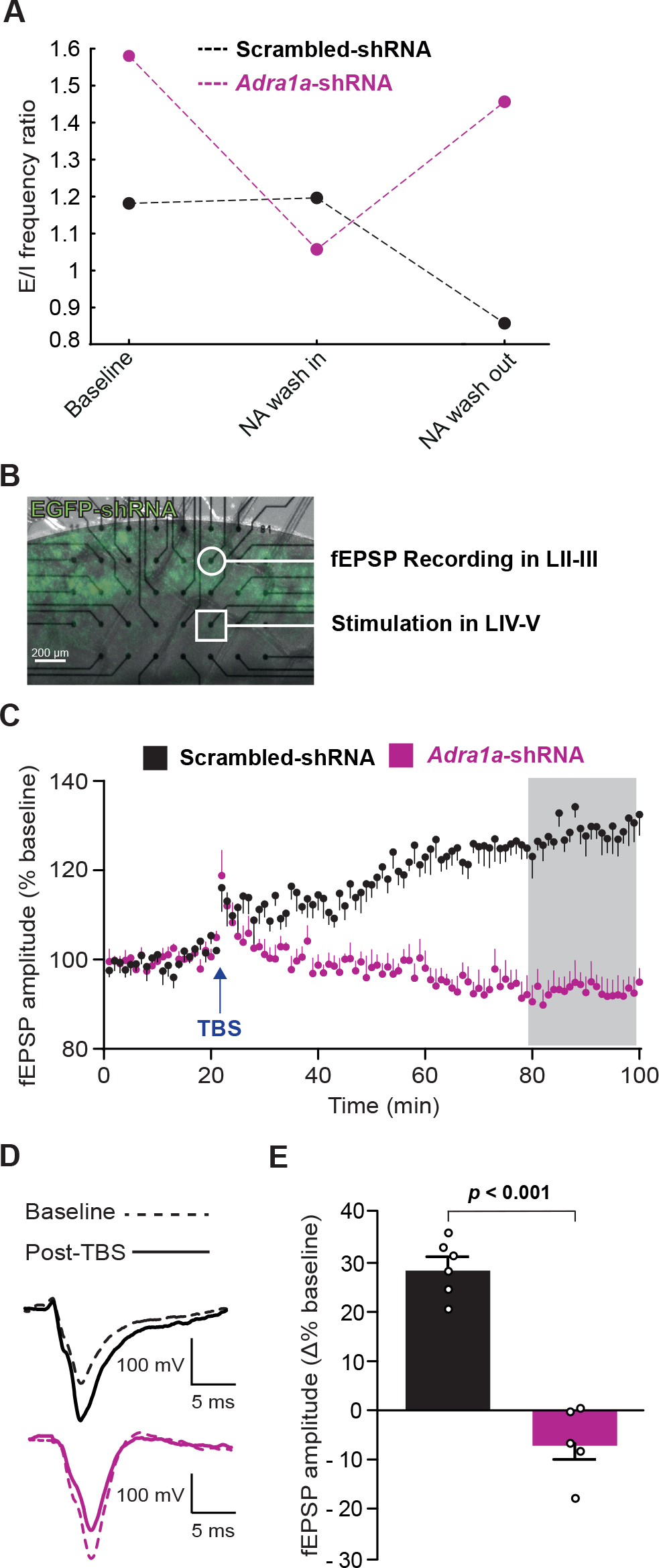
Astrocyte α1A-NAR is crucial for LTP induction in V1 A. This graph visually represents the ratios between the average E- and I-PSC frequency values as measured in Figure 4E, G (barplots, right) for baseline, NA wash in and NA wash out conditions. B. Example image of an acute brain slice of V1 composed of a brightfield image of the slice positioned on an MEA chip and overlaid with an IHC image of the same slice revealing EGFP-shRNA expression in layer II- III. The white circle and square represent the typical positions of electrodes chosen to acquire or evoke an fEPSP, respectively. Only slices in which there was visible EGFP expression in layers II-III were used. C. Time course of the averaged fEPSP amplitudes relative to the baseline average. The point at which theta burst stimulation (TBS) was applied is indicated with a blue arrow. Vertical bars represent the SEM. The grey box represents the timepoints averaged to produce the plots in (E). D. Example traces of the evoked fEPSP for two recordings from each experimental group; dashed traces represent the fEPSP during baseline, solid traces the fEPSP after TBS. E. Barplots of the variation of fEPSP amplitude averaged per group over all timepoints between the 79th and 99th minutes of the recording (highlighted by the grey box in (C)). Circles indicate the fEPSP amplitude averaged per slice, error bars represent the SEM. Scrambled-shRNA: n = 3 mice; 6 slices. Adra1a-shRNA: n = 3 mice; 5 slices. Statistic above the graph in (E) indicates the p-value for an independent sample t-test. All descriptive statistics and statistical test parameters are summarized in Table S1.

### Preparation of acute brain slices

Preparation followed a published protocol (Ting et al., 2018) and was performed as follows. Animals were anesthetized using intraperitoneal administration of nembutal (50 mg/kg). Transcardial perfusion was performed using 20 ml of ice-cold N-methyl-D-glucamine-based artificial cerebrospinal fluid (NMDG-ACSF) dissection solution, containing (in mM): NMDG 93, KCl 2.5, NaH_2_PO_4_ 1.25, NaHCO_3_ 30, MgSO_4_ 10, CaCl_2_ 0.5, HEPES 20, D-glucose 25, L-ascorbic acid 5, thiourea 2, sodium pyruvate 3, N-acetyl-L-cysteine 10; pH 7.4 (HCl). Osmolarity was adjusted to 305–310 mOsm/l, if needed. The solution was bubbled in 95% O_2_/5% CO_2_ gas for 20 min before use and bubbling was maintained throughout the experiment.

Following perfusion and decapitation, the brain was swiftly removed. Coronal slices, containing the visual cortex (350 µm-thick), were obtained using a VT1200s vibratome (Leica Biosystems). Once obtained, slices were further cut along the midline in ice-cold NMDG-ACSF using a scalpel blade to obtain 2 independent sections corresponding to the left and right brain hemisphere and placed in a chamber containing NMDG-ACSF at 33°C. Slices were maintained under these conditions for 25 min, with the controlled reintroduction of Na^+^ achieved by gradual addition of 2 M NaCl to the chamber, based on the published protocol (Ting et al., 2018). For patch-clamp and Ca^2+^ imaging experiments, slices were then transferred to another chamber containing room temperature recording ACSF (in mM): NaCl 124, KCl 2.5, NaH_2_PO_4_ 1.25, NaHCO_3_ 26, MgSO_4_ 2, CaCl_2_ 2, D-glucose 15; pH 7.4 (HCl). For MEA recordings, we used a modified recording ACSF (in mM): NaCl 124, KCl 4.5, NaH_2_PO_4_ 1.25, NaHCO_3_ 26, MgCl_2_ 1, CaCl_2_ 2.5, D-glucose 10, pH 7.4 (HCl). ACSF solutions were routinely bubbled with a 95% O_2_/5% CO_2_ gas mixture and had an osmolarity in the range of 305–310 mOsm/l. For calcium imaging experiments, slices were removed from the storage chamber and transferred to a six-well plate for loading with Rhod-2-AM (see below), before being returned to the storage chamber. For calcium imaging and electrophysiological recordings, slices were transferred to a specialized recording chamber and superfused with recording ACSF at 22°C (patch-clamp and Ca^2+^ imaging experiments), or modified recording ACSF at 32°C (MEA experiments), including pharmacological reagents, where appropriate.

### Ca^2+^ imaging

Dye loading and 2-photon microscopy: Following brain slicing, slices containing V1 were incubated in a solution containing Rhod-2-AM. The dye was supplied as a 50 µg ampoule (Thermo Fisher Scientific) and was solubilized using a mixture of 7 μl dimethyl sulfoxide (DMSO), 2 μl 20% Pluronic F-127 in DMSO (Tocris Biosciences), and 1 μl 0.5% Kolliphor EL (Sigma-Aldrich) in DMSO. The ampoule was then incubated at 41°C with constant agitation (1400 RPM) for 15 min using a thermomixer. Concentrated Rhod-2-AM was then added to a well containing 3 ml of recording ACSF, giving a final concentration of 14.8 µM. Rhod-2-AM was chosen for its red-shifted emission profile, allowing us to simultaneously image the EGFP fluorescence with minimal spectral overlap. Slices were loaded in this solution for 45–60 min at 35°C. At the end of this period, excess AM dye was removed by washing in room temperature recording ACSF for at least 1 h. Prior to use, slices were maintained in a storage chamber containing room temperature recording ACSF.

All Ca^2+^ imaging experiments followed the same protocol. A field of view containing layers II-III of the primary visual cortex containing EGFP-shRNA expressing astrocytes was chosen. This field of view was then recorded under two sequential conditions. First, recording ACSF containing 50 µM (R)-(-)- Phenylephrine hydrochloride, (PHE, Tocris Bioscience) was applied at t + 60 s and maintained for the remaining duration of the recording period (240 s). After washing the slice in recording ACSF for 900 s, we then repeated the protocol adding 10 µM ionomycin (Sigma-Aldrich) at t + 60 s. This Ca^2+^ selective ionophore was chosen to induce a positive response to control for astrocyte viability in the slice (see “Ca imaging data analysis“ below).

Live imaging of cells in acute slices was performed using a two-photon imaging system (VIVO 2-Photon platform, Intelligent Imaging Innovations GmbH), equipped with a tunable multiphoton laser (MaiTai laser, Spectra-Physics). Imaging was performed using a Zeiss Axio Examiner Z1, equipped with a W PlanApochromat 20×/NA 1.0 objective. To excite both EGFP and Rhod-2, the excitation wavelength was tuned to 900 nm. Fluorescence signals were split using a 580 nm edge BrightLine single-edge imaging-flat dichroic beam splitter (Semrock FF580-FDi01-28x38) and detected using two fast-gated GaAsP PMTs (Hamamatsu Photosensor Modules H11706) equipped with either a 525/40 nm or a 612/69 nm emission filter. Images were 512 × 512 pixels in size and acquired at a frequency between 2 and 4 Hz. Acquisition was controlled using Slidebook 6 software (Intelligent Imaging Innovations GmbH). Laser power was limited to a maximum of 30 mW at the specimen. The focal plane used was usually 30–100 μm within the slice.

Ca imaging data analysis: Images were initially processed using Fiji (Schindelin et al., 2012) with standard plugins to correct for image drift and noise. ROIs encompassing the cell soma were superimposed onto the images of visually identified EGFP-shRNA positive astrocytes. Where possible, multiple ROIs per image were used. First, the ‘sum slices’ projection from the Fiji ‘Z stack’ plugin was applied to all frames acquired in the EGFP channel, to evaluate the EGFP intensity in each ROI. Next, the average pixel intensity per frame of the Rhod-2 channel was then measured for each ROI and exported to MATLAB (The Mathworks). Following this, we used custom-written scripts to evaluate the relative variations in [Ca^2+^], estimated as changes in Rhod-2 signal relative to the baseline (essentially ΔF/F_0_). The baseline (F_0_) was defined for each ROI as the average fluorescence across frames before drugs were added to the recording chamber. To avoid negative ΔF/F_0_ values, when ΔF was slightly negative, we added 1 to the calculation of the ΔF/F_0_ for each individual frame. The ΔF/F_0_ ratios were then converted into percentages, with a value of 100% equivalent to an absence of change relative to baseline. Only astrocytes that demonstrated a ΔF/F_0_ amplitude greater than 110% following ionomycin application were kept for further analysis. Of these astrocytes, we calculated the amplitude of the response to PHE, the time needed for the maximal PHE response to occur (time to peak), as well as the total area under the curve (AUC) of the ΔF/F_0_ trace.

### Electrophysiology

Whole-cell patch-clamp recordings: Recordings were done using borosilicate glass recording electrodes (with resistances typically between 3.5–5 MΩ). We used the following internal solution (in mM): 115 CsMSF, 20 CsCl, 10 HEPES, 2.5 MgCl_2_, 4 ATP, 0.4 GTP, 10 creatine phosphate, 0.6 EGTA and 0.2% biocytin (pH 7.25), with osmolarity adjusted to 290–300 mOsm/l, if needed. Recordings were performed using a Multiclamp 700B amplifier (Axon Instruments). Series capacitances and resistances were compensated electronically throughout the experiments using the main amplifier. To isolate inhibitory post-synaptic currents (IPSCs), neurons were clamped at a holding potential of 0 mV. To isolate excitatory post- synaptic currents (EPSCs), neurons were clamped at a holding potential of -70 mV. The recordings were conducted as follows: first, spontaneous post-synaptic currents (PSCs) were recorded for 360 s. At t + 360 s, the chamber was perfused with recording ACSF containing 20 µM noradrenaline bitartrate (Tocris Bioscience) for 60 s. Next, the drug was washed from the chamber using standard recording ACSF and the recording continued until t + 900 s. The amplitude and frequency of post-synaptic currents (PSCs) were analyzed using the Mini Analysis program (Synaptosoft), in a semi-automated fashion (automatic detection of events with chosen parameters followed by visual inspection and validation). Further analyses were performed using MATLAB (The Mathworks).

MEA recordings: These recordings were performed using modified recording ACSF. The MEA setup was composed of a Multichannel Systems (MCS) MEA2100-Mini-Headstage coupled to a MCS interface board 3.0 multiboot, connected to a MCS 60MEA200/30iR-Ti MEA chip composed of 60 electrodes uniformly spaced by 200 µm. The set-up was also equipped with a TC02 temperature regulator and a PPS2 peristaltic pump (both supplied by MCS). Acute brain slices were selected for recordings based on the presence of EGFP-shRNA fluorescence in V1. The slice was then positioned on the MEA chip in an orientation enabling stimulation of neurons in cortical layers IV-V and recording of evoked potentials in cortical layers II-III. Slices were maintained firmly in contact with the MEA chip using a custom-made mesh of nylon wires tied to a stainless steel ring, which was positioned over the slice. Following a stabilization period of 45 minutes following placement onto the MEA chip, slices were submitted to an input-output (IO) protocol. Each stimulus in the IO protocol consisted of three pulses of one given amplitude, starting from 500 mV. Stimuli were repeated every 10 seconds with incremental increases in amplitude of 250 mV, until a final amplitude of 3000 mV was reached. We then selected the stimulus strength which evoked a field excitatory post-synaptic potential (fEPSP) corresponding to 35% of the maximal amplitude reached by the fEPSP in the IO protocol. This stimulus strength was then used throughout the recording, including during theta-burst stimulation (TBS). Our recording protocol consisted of a 100 minutes long recording, with 1 stimulus per minute. At t + 21 min, we applied a TBS, composed as follows: an episode of 5 stimulations was applied at 100 Hz and repeated 50 times (with a 200 ms interval between each episode). All pulses used were biphasic with a pulse width of 600 µs. fEPSP amplitudes values were extracted using MCS analyzer software. These data were subsequently fed into a MATLAB script. The script computed the change in amplitude relative to the average baseline (fEPSP amplitudes between t + 0 and t + 20 minutes) expressed as a percentage, with baseline set to 100%. Values of fEPSPs elicited during the TBS stimulation were excluded from analysis. We excluded recordings in which the standard deviation of the baseline values exceeded 6%.

### Noradrenergic receptors expression profile analysis

Analysis was performed on a previously published single cell RNA-seq database (Batiuk et al., 2020), using Seurat version 4.1.0, run on RStudio version 1.3.1093, using R version 4.0.3. 990. Cortical astrocytes (AST2 and AST3 in the dataset) were identified and the associated expression data used for downstream analysis. Data were log-normalized using the default normalization method in Seurat (Hao et al., 2021). The figure was generated with R ggplot2 package version 3.4.2. Data distribution normality was checked using a Shapiro-Wilk test, using Stats version 3.6.2. Kruskal-Wallis and Dunn’s tests were performed with R rstatix package version 0.7.2; Spearman correlation test was performed with ggpubr version 0.6.0.

### In vitro Immunoassays

HEK293T cells were maintained in DMEM supplemented with 10% FBS and 1% penicillin – streptomycin (complete medium) and maintained in an incubator (37°C, 5% CO_2_).

Transduction with lentivirus: To express EGFP-shRNA, HEK293T cells were transduced overnight with a single lentiviral vector expressing the three Adra1a-targeting shRNAs, or with a lentiviral vector expressing the scrambled (control) shRNA-expressing. In both cases, shRNA expression was under the control of the ubiquitous CMV promoter. The shRNA sequences used were the same as those described above (see “AAV-mediated, astrocyte-specific shRNA expression strategy”). Vectors used in different experiments were all from the same batch, and produced by VectorBuilder. For transduction, medium was changed to complete DMEM containing lentivirus (multiplicity of infection: 5) and 5 μg/ml polybrene. As a negative control, cells were treated only with polybrene (non-transduced). 5 days after transduction, cells were split into either 6 well-plates (300000 cells per well) coated with 1 mg/ml Poly-L-Lysine (PLL, Merck, P9155) for western blotting, or into 24 well-plates (95000 cells per well) with PLL-coated coverslips for immunofluorescence.

Transfection with α1A-V5(x2)-NAR: The next day, cells were transfected with a plasmid, based on the mCherry2-N1 backbone. The plasmid encoded a recombinant receptor consisting of an α1A-NAR fused to two V5 epitope tags at the extreme N-terminus of the protein (α1A-V5(x2)-NAR), separated from mCherry (transfection marker) using a P2A sequence. Expression of α1A-V5(x2)-NAR was driven by a CMV promoter for ubiquitous expression. Transfection was performed using polyethylenimine (PEI) at a stock concentration of 1 mg/ml (in sterile water). For each well of a 6 well-plate, 2.67 μg of plasmid DNA diluted in 340 μl of a 150 mM NaCl solution was mixed with 9.34 μg PEI stock diluted in 340 μl of a 150 mM NaCl solution. For each well of a 24 well-plate, 0.53 μg of plasmid DNA diluted in 100 μl of a 150 mM NaCl solution was mixed with 1.87 μg PEI stock diluted in 100 μl of a 150 mM NaCl solution. These solutions were then mixed and incubated for 20 minutes at room temperature. Subsequently, each PEI/DNA solution was further mixed with DMEM containing 1% FBS to obtain the transfection medium (3 ml per well of a 6 well-plate and 500 μl per well of a 24 well-plate). Complete cell culture medium was replaced with transfection medium and cells were incubated for 5 hours (37°C, 5% CO_2_). After 5 hours of incubation, transfection medium was removed from the cells and complete cell culture medium was added. This protocol (transduction followed by transfection) was performed three different times to obtain three biological replicates.

Immunoblotting: protein lysates were prepared 3 days after transfection by collecting cells directly into ice-cold RIPA buffer (50 mM Tris-HCl pH 7.5, 150 mM NaCl, 1% Triton X-100, 1% Na-Deoxycholate, 0.1% SDS) supplemented with a protease inhibitor cocktail (Roche, 4693159001). Samples were centrifuged at 15700 g, for 20 minutes at 4°C and the supernatant was collected into a 1.5 ml centrifuge tube. Protein concentration was determined with a BCA Protein Assay Kit (Thermo Scientific, 23225), using Bovine Serum Albumin (BSA) as a standard. 10 μg of the protein samples were mixed with 4X Laemmli Sample Buffer (BioRad, 1610747) containing 2-mercaptoethanol. Samples were then boiled at 95°C for 5 minutes before loading onto the gel. PageRuler™ Prestained Protein Ladder (ThermoScientific, 26616) was used as a ladder to determine the molecular weight of protein samples. Samples were run on an SDS-PAGE gel (5% stacking gel and 10% separating gel) at 100 V for 120 minutes, in 1X SDS Running Buffer, pH 8.3. Proteins were transferred to a methanol-activated Immuno-Blot PVDF Membrane (BioRad, 1620177) using tank-transfer at 300 mA, for 1.5 hours at 4°C. Following transfer, the membrane was incubated with Ponceau S for 5 minutes to ensure successful protein transfer and then incubated in a blocking buffer (5% Non-fat Dried Milk Powder in TBS-T (0.1% Tween20, Tris-buffered saline) for 1 hour at room temperature. The membrane was next incubated with primary antibody diluted in 2% BSA in TBS-T at 4°C overnight. Next day, the membrane was rinsed three times with TBS-T, with each wash lasting 5 minutes. Membranes were incubated with a secondary antibody diluted in blocking buffer at room temperature for 1 hour. Then, the membrane was washed three times with TBS-T, with each wash lasting 5 minutes. Proteins were detected using Pierce ECL Western Blotting Substrate (Thermo Scientific, 32109) and visualized using an iBright 1500 imaging system (Invitrogen). As expected, based on predicted molecular weight, α1A-V5(x2)-NAR was detected by a major band at approximately 60 kDa molecular weight. Quantification was performed by measuring the intensity of this V5 band, as well as the intensity of the vinculin band using the ‘Gel Analysis’ Fiji plugin (Schindelin et al., 2012). For each condition, V5 band intensity was normalized to its respective vinculin loading control using Microsoft Excel. Antibodies and dilutions used were as follows: mouse anti-V5 (Invitrogen, R960-25, 1:1000), mouse anti-vinculin (Sigma, V9131, 1:2000) and goat anti-mouse HRP-conjugate (BioRad, 1706516, 1:1000).

Immunofluorescence: 3 days after transfection, cells were fixed with 4% paraformaldehyde solution in PBS for 10 min and then washed with three times in PBS, with each wash lasting 5 minutes. Cells were incubated in blocking solution (3% BSA + 0.3% Triton X-100 in 1X PBS + 0.02% Na-azide) for 1 hour at room temperature. After blocking, cells were incubated in primary antibodies, diluted in blocking solution, overnight at 4°C. Next day, cells were washed with PBS (3 washes each of 5 minutes) and incubated with secondary antibodies diluted in blocking solution for 1 hour at room temperature. After labelling with secondary antibodies, cells were washed with PBS (3 x 5 minutes) and mounted on standard microscope glass slides using DAPI-Fluoromount G (SouthernBiotech, 0100-20). Antibodies and dilutions used were as follows: chicken anti-GFP (1:5000 dilution, Aves Labs, GFP-1010); RFP booster-594 (1:400 dilution, ChromoTek ATTO-594, rba594), mouse anti-V5 (1:500, Invitrogen, R960-25), Alexa Fluor- 488 donkey anti-chicken (1:1000, Jackson ImmunoResearch, 703-545-155) and Cy5-donkey anti-mouse (1:1000, Jackson ImmunoResearch, 715-175-150). Cells were imaged using an inverted Nikon TiE A1R confocal microscope equipped with a W PlanApochromat 20×/NA 1.0 objective. For analysis, ROI were drawn automatically (using the mCherry signal) and the expression of α1A-NAR (based on the intensity of V5-epitope tag staining) extracted using QuPath (Bankhead et al., 2017). Further analyses were performed using MATLAB (The Mathworks) and plots obtained using Prism (GraphPad Software).

### Experimental design and statistical analysis

Data are presented as the mean +/- standard error of the mean (SEM), unless otherwise stated. Statistical tests were chosen according to the distribution of the data and are indicated in the figure legends, as well as in Table S1. Tests were performed using either MATLAB (The Mathworks), Prism (GraphPad Software) or R ggsignif package version 0.6.2.

## RESULTS

We initially used our previously generated scRNA-seq astrocyte database to analyze the expression profile of the nine known mouse NAR genes (Figure 1). This revealed that Adra1a is the major transcript found in cortical astrocytes, both in terms of the number of cells positive for this transcript and the average expression level in each cell. Based on this transcriptome data, therefore, we concentrated our experiments on Adra1a. Unfortunately, a floxed mouse line allowing astrocyte-specific Adra1a cKO was not available to us at the time we initiated this study. Given this fact, we chose to use a short hairpin RNA (shRNA)-based approach for flexible, astrocyte-specific KD (Fellmann et al., 2013). We identified three miR-30-based shRNAs that targeted the Adra1a in a non-overlapping manner and decided to use all three sequences in combination to obtain the best possible gene KD. Prior to experiments in mice, we tested the efficiency of our strategy using a simple protocol based on overexpression of an epitope-tagged α1A-NAR in HEK293T cells (Figure S1A). We tested the ability of our chosen shRNA sequences to decrease α1A-NAR using both immunocytochemistry (Figure S1B) and Western blotting (Figure S1C). Both approaches showed that our chosen shRNA sequences reduced α1A-NAR levels (compared to a scrambled control). Knowing our KD strategy worked in vitro, we next sought to express the same shRNAs in V1 adult mouse astrocytes. We took advantage of a commercially available FLEx system (Schnütgen et al., 2003), delivered using an adeno-associated virus (AAV)-based vector system, to avoid major issues with astrocyte reactivity (Chatterjee et al., 2021). The FLEx system allows Cre-mediated switching between expression of two open reading frames within a single expression cassette. In our system, transduced cells expressed mStrawberry in the absence of Cre activity. Cre-mediated recombination was used to turn on shRNA expression, with EGFP co-expression serving as a recombination marker. We obtained three independent AAV vectors, each encoding one of the Adra1a- targeting shRNA we previously identified. The three AAV solutions were mixed in roughly equal amounts prior to stereotactic injection. As a control, we used an AAV with the same scaffold, but expressing a scrambled shRNA (Figure 2A). When referring to vectors without distinguishing between the shRNAs expressed, we use the term “EGFP-shRNA”. When referring to the AAV mixture targeting Adra1a, we use the terminology “Adra1a-shRNA”. Finally, when referring to the control vector, we use “scrambled-shRNA”. To obtain astrocyte-specific expression of the EGFP-shRNA constructs, we injected these AAVs into V1 of Fgfr3-iCreER^T2^ transgenic mice (Young et al., 2010), which allows for highly efficient and specific recombination in cortical astrocytes following tamoxifen administration (Hu et al., 2020) (Figure 2A). This acute KD strategy allowed us to control both the timing and specificity of Adra1a-shRNA expression, avoiding potential developmental issues, possible genetic compensation and/or the lack of cell type specificity occasionally seen when using AAV vectors in combination with supposedly astrocyte-specific promoter systems (O’Carroll et al., 2021 and references therein).

To ensure the cell-type specificity of our KD, which is crucial for our experiments, we first used immunohistochemistry (IHC) to determine the cell type specificity of EGFP (and hence shRNA) expression in the supragranular V1. We found EGFP was expressed in over 35% of S100β positive astrocytes, and (importantly) less than 0.2% of NeuN positive neurons (Figure 2B). This indicates that while our strategy had only moderate efficiency in terms of cell coverage, it was highly specific for astrocytes.

Knowing we could target EGFP-shRNA expression correctly, we next sought to test if the expression of Adra1a-targeting shRNA modified astrocyte responses following exposure to an α1-NAR agonist. To this end, we measured changes in [Ca^2+^] in Adra1a-shRNA or scrambled-shRNA positive astrocytes using the red-shifted Ca^2+^ indicator Rhod-2 (Figure 3A), which has been previously used to monitor calcium transients in astrocytes, including after α1-NAR activation (Tran et al., 2018; Institoris et al., 2021). We activated α1-NAR signaling using the pan-α1-NAR agonist, phenylephrine (PHE). We normalized the change in Rhod-2 fluorescence over time to the averaged baseline value before PHE application (essentially ΔF/F_0_, Figure 3A-B). While most recorded astrocytes still displayed increases in [Ca^2+^] following PHE application (Figure 3), we found a positive correlation between the fluorescence intensity of the EGFP reporter and the time at which the [Ca^2+^] response peaked in the Adra1a-shRNA group (coefficient estimate: 3.3259, p-value: 4.0059 x 10^-5^, R-squared: 0.156, Figure 3C), which was absent in the scrambled-shRNA group (coefficient estimate: -1.3153, p-value: 0.53296, R-squared: 0.00459, Figure 3C). This is exemplified by the traces displayed in Figure 3B, where the astrocyte with the lowest EGFP intensity (cell 1 in Figure 3A-B) has the fastest and highest increase in [Ca^2+^], whereas cell 3, which shows the highest EGFP intensity, has a delayed and diminished response to PHE. Following this observation, we decided to bin our measurements depending on the measured EGFP fluorescence, based on the assumption that EGFP intensity is a proxy for the level of shRNA expression and hence Adra1a KD. Similar dose-dependent effects of shRNA expression on KD efficacy have already been reported for shRNA targeting various mRNAs, both in vitro (Gupta et al., 2004; Suhy et al., 2012) and in vivo (Koornneef et al., 2011; Wang et al., 2012). For further analysis, we segregated the astrocytes into three equally sized bins, based on EGFP intensity. The ‘low’ EGFP intensity group was composed of astrocytes whose EGFP intensities were in the lowest third of those measured; the second ‘high’ intensity group gathered astrocytes belonging to the two other bins (dashed line in Figure 3C, low EGFP intensity versus high EFGP intensity cells in Figure 3D-E). We next quantified the [Ca^2+^] changes by computing the amplitude, time to peak and area under the curve (AUC) for the ΔF/F_0_ trace of each cell. While in the low EGFP intensity group there was no differences in the averaged amplitude or AUC of the ΔF/F_0_ traces between scrambled- or Adra1a-shRNA groups, the averaged time to peak was already significantly longer in the Adra1a-shRNA group (Figure 3D). In the group composed of high EGFP intensity astrocytes, we found that the averaged response to PHE was significantly lower in amplitude, had a longer time to peak and, consequently, had a lower AUC in the Adra1a-shRNA group compared to the scrambled-shRNA group (Figure 3E). This indicated to us that while not completely knocking down the response of astrocytes to α1-NAR activation by PHE, our KD strategy clearly altered it.

Based on this effect, we decided to continue our experiments using this Adra1a KD strategy. In order to assess the specific contribution of astroglial α1A-NAR to the complete noradrenergic response in V1 circuits, we bath applied NA and measured the effects on synaptic inputs to visually identified pyramidal neurons located in layers II-III of V1. To do this, we patched neurons in an area where EGFP-expression in astrocytes was clearly visible, and clamped the neurons at membrane potentials that allow either excitatory or inhibitory post-synaptic currents (E- or I-PSCs) to be measured (Figure 4A-C). We measured the baseline frequency and amplitude of the PSCs, then bath-applied NA for 60 s and finally washed it out for the remainder of the recording (Figure 4A-D). First, we quantified the measured PSCs by binning the averaged frequency and amplitude values for all recorded neurons, across each experimental group, using 30 s bins spanning the entire recording period (Figure 4E-H, left). Next, we adopted a similar strategy but averaged the measured PSC parameters in defined 60 s bins (as this corresponds to the duration of NA bath-application) (Figure 4E-H, right). We found that the α1A-NAR KD in astrocytes affected IPSC frequency during baseline and NA wash out, but not the increase in IPSC frequency measured during NA wash in (Figure 4E). Moreover, we found a slight but significative increase of IPSC amplitude during NA application in the scrambled-shRNA group, an effect lost in the Adra1a-shRNA group (Figure 4F). Regarding EPSCs, we observed that while there was a non-significant trend towards increased EPSC frequency during NA wash in for both groups (Figure 4G), no statistically significant differences could be found between the two EGFP-shRNA-expressing groups (Figure 4G-H).

Based on these results, we plotted the ratio of EPSC over IPSC frequencies during baseline, NA wash in and NA wash out (Figure 5A) to estimate an E/I ratio (Zhou et al., 2009; Bassetti et al., 2021; Kirischuk, 2022). This plot highlights that in the Adra1a-shRNA group, there is an imbalance in favor of excitation, when the circuit is either in its basal state, or following the wash out of NA from the recording bath. We reasoned that such an imbalance could alter the capacity of circuits in layers II-III of V1 to undergo plasticity. Indeed, reduced inhibitory tone has been linked to impairments in the induction of long-term potentiation (LTP) following high frequency stimulation (Jedlicka et al., 2009), with one study even reporting a switch to long term depression (LTD) (Le Roux et al., 2008). Furthermore, α1-NAR signaling and astrocytes have each been independently implicated in LTP induction in circuits spanning layers IV-VI to layers II-III of V1 (Pankratov and Lalo, 2015). However, a direct link between these two observations has, to date, not been reported. Therefore, we tested whether astrocyte-specific α1A-NAR KD resulted in reduced plasticity in V1. To do this, we used a multi-electrode array (MEA) system to simultaneously stimulate neurons in layers IV-V and record the evoked field excitatory post-synaptic potentials (fEPSP) in layers II-III (Figure 5B). We found that application of a tetanic theta-burst stimulation (TBS) protocol led to LTP in the control (scrambled-shRNA) group. The same protocol applied in the Adra1a-shRNA group revealed that not only was LTP lost, but that LTD was actually induced (Figure 5C-D). These results demonstrate that α1A-NAR signaling in astrocytes is crucial for normal circuit function, notably plasticity, in V1.

## DISCUSSION

In this study, we confirm that NA signaling in V1 astrocytes relies on the α1A-NAR subtype (Figures 1 & 3). We provide evidence that alterations in α1A-NAR signaling in astrocytes affect inhibitory synaptic transmission in layers II-III of V1 (Figure 4), tipping the E/I balance of the circuit in favor of excitation, with drastic consequences for synaptic plasticity in this area (Figure 5).

Our results provide valuable insights, but also raise new questions about α1-NAR signaling in V1. To a degree, these questions arise because of the methodological approach adopted. At the time we initiated the project, there were no floxed Adra1a mouse lines readily available, which is why we adopted a shRNA-based approach. One of the potential limitations of shRNA is variable (dose-dependent), and possibly incomplete, gene KD (Gupta et al., 2004; Koornneef et al., 2011; Suhy et al., 2012; Wang et al., 2012; Boettcher and McManus, 2015). Indeed, our in vitro experiments testing the efficiency of α1A-NAR KD (Figure S1) suggests that this is likely. However, it should be pointed out that this experiment may actually underestimate receptor KD in astrocytes for two main reasons. First, our strategy was based on the overexpression of an epitope-tagged α1A-NAR in HEK cells (to allow ease of detection), at levels which are likely to exceed the endogenous amounts expressed in adult mouse astrocytes. Second, not all HEK cells overexpressing the tagged α1A-NAR were transduced with the Adra1a-shRNA, which would artificially reduce the average KD efficacy we measured using simple Western blot and immunocytochemistry approaches. However, incomplete receptor KD is, in our opinion, the most parsimonious explanation for what appears to be a partial loss of astrocyte responses to α1-NAR activation in the Adra1a-shRNA group (Figure 3).

Variable levels of receptor KD would also explain the apparent dose-dependent effects we observed, with astrocytes expressing low levels of Adra1a-shRNA displaying only slight alterations in their [Ca^2+^] responses, whereas cells with comparatively higher levels of Adra1a-shRNA expression displayed a heavily modified [Ca^2+^] increase following PHE application, with drastically altered kinetics and amplitude (Figure 3). While we acknowledge that the observed correlation between EGFP fluorescence and time to peak ΔF/F_0_ (Figure 3C) is not especially strong (R-squared: 0.156), we still consider this the best explanation for the data. Furthermore, a recent study, in which the authors employed Adra1a cKO transgenic mice, but also injected a cohort of animals with an AAV expressing Adra1a-targeting shRNA in astrocytes, further supports this hypothesis. Indeed, Kohro and colleagues found a significant but incomplete loss of astrocyte responses to PHE following Adra1a-targeting shRNA expression, similar to our results, while these responses were much lower to absent in the cKO mice. Crucially, however, these authors found similar behavioral phenotypes between the KD or cKO mice, highlighting that even partial KD of α1A-NAR in astrocytes can be sufficient to produce a clear phenotype by reducing the effect of α1- NAR activation on circuits responsible for pain perception and response (Kohro et al., 2020). Interestingly, our study reaches a similar conclusion, with reduction of astrocyte [Ca^2+^] signaling, through partial α1-NAR knock down, sufficient to produce clear-cut effects on cortical circuit activity (Figure 5).

Despite the attractiveness of this hypothesis, however, we cannot formally exclude the possibility that a residual degree of Ca^2+^ signaling in astrocytes is mediated by other NARs. Based on our single cell sequencing data, cortical astrocytes also express Adra1b, Adra2a and Adrb1. These encode for the α1B- NAR, α2A-NAR and β1-NAR, respectively. Some reports indicate that agonists for β-NAR can induce [Ca^2+^] transients in astrocytes in vivo (Bekar et al., 2008) and in vitro (Horvat et al., 2016) while one study reported α2-NAR activation evoked [Ca^2+^] transients in a minority of cultured astrocytes (Gaidin et al., 2020). However, we feel it unlikely that α2A-NAR and β1-NAR play significant roles in the residual [Ca^2+^] response observed here (Figure 3), based on the fact that (i) their expression is limited to comparatively few cells (which surprisingly do not appear to co-express α1A-NAR, Figure 1 and Table S2) and (ii) in vivo studies which show that cortical astrocytes [Ca^2+^] responses to phasic NA release are fully prevented by α1-NAR antagonists, while β- or α2-NAR blockers have no effects (Ding et al., 2013; Oe et al., 2020). Moreover, since we used an α1-NAR selective agonist, PHE, α2A-NAR and β1-NAR activation would need to be triggered by endogenous NA retained in our brain slice preparation. Based on these arguments, we believe that outside an incomplete KD, the only possible explanation to the residual degree of Ca^2+^ signaling observed is stimulation of α1B-NAR signaling by PHE. This interpretation is consistent with the findings of recent studies based on genetic ablation of α1A-NAR. Using either a constitutive (global) Adra1a KO, or a recently generated line with a floxed Adra1a allele (for cell-type specific KO), the Paukert lab reported that functional ablation of α1A-NAR expression in Bergmann glia resulted in a major, but not complete, loss of NA-induced Ca^2+^ transients in these cells, with the transient dynamics (reductions in amplitude and time to peak), similar to the observations we report here. Since both bulk and scRNA-seq data indicate that the Adra1b transcript is present in significant amounts in cerebellar astrocytes (Boisvert et al., 2018; Saunders et al., 2018), it is possible that this ‘residual’ response to NA results from α1B-NAR activation. Likewise, a study exploring the effects of Adra1a KO in a molecularly defined subtype of astrocytes (Hes5), located in the superficial laminae of the mouse dorsal horn, reported a near complete loss of [Ca^2+^] increases (Kohro et al., 2020). While several reports claim that both Adra1a and Adra1b are expressed in spinal cord astrocytes, (Anderson et al., 2016; Blum et al., 2021; Russ et al., 2021), studies using scRNA-seq approaches seem to indicate this only holds for a subset of astrocytes (Blum et al., 2021; Russ et al., 2021). Interestingly, when examining our own single cell data, we also found a similar expression pattern in cortical astrocytes, with only six astrocytes co-expressing both genes (Figure 1, Table S2), suggesting that Adra1a and Adra1b positive astrocytes may actually represent two distinct subpopulations in the cortex. Hence, it remains unclear to what degree α1A-NAR and α1B-NAR play roles in astrocyte Ca^2+^ signaling. We believe that the cleanest way to disentangle the relative contributions of both receptor types to α1-NAR signaling in V1 would be to replicate our experiments in mice in which both Adra1a and Adra1b genes are KO in astrocytes, both individually and in combination, to assess the effects on astrocytic Ca^2+^ increase following α1-NAR activation. Whether any such modification would further exert an effect on synaptic function and plasticity is difficult to predict: despite being G_q_-coupled like α1A-NAR, it is possible that α1B-NAR actually activates different signaling pathways downstream, or in parallel, to increases in [Ca^2+^] (Cotecchia, 2010). This hypothesis is supported by studies showing that mice overexpressing constitutively active forms of α1A- or α1B-NAR display drastically different phenotypes, such as lifespan (Collette et al., 2014). Such differences between α1A- and α1B-NAR signaling pathways downstream or parallel to an otherwise common effect on [Ca^2+^] could also explain the apparent discrepancy between our observation of residual increases in astrocyte [Ca^2+^] following PHE application (Figure 3) and the rather clear-cut effects observed on synaptic function and plasticity in mice with astrocytic KD of α1A- NAR (Figures 4 and 5).

However, despite these caveats, it is still remarkable that even a partial/incomplete α1A-NAR KD (Figure S1), in a limited proportion of layer II-III astrocytes (35%, Figure 2) is sufficient to exert profound effects at the synaptic level, with a reduction in IPSC frequency measured during both baseline and NA wash out, and a moderate loss of increase in IPSC amplitude seen during NA wash in (Figure 4E-F).

Interestingly, we observed somewhat similar results in our study regarding the role of astrocytes in oxytocinergic signaling, with the loss of oxytocin receptor expression in a small subpopulation of astrocytes being sufficient to drastically impair the effect of oxytocinergic signaling on inhibitory transmission in central amygdala circuits (Wahis et al., 2021a). In the visual cortex and hippocampus, astrocytes have an essential role in maintaining the inhibitory tone on pyramidal neurons during firing of somatostatin (SST) positive interneurons (Matos et al., 2018; Henriques et al., 2022), a role which parallels our observation that NA-induced increases in inhibitory current amplitude and frequency are diminished and shortened, respectively, by α1A-NAR KD in astrocytes (Figure 4E). Mechanistically, this control of basal GABAergic transmission by astrocytes could rely on the regulation of a number of known astrocyte functions. For example, astrocytes control neuronal excitability by controlling levels of extracellular neurotransmitters (including GABA) (Liu et al., 2022) and extracellular K^+^ levels. Moreover, these cells also play essential roles in (inhibitory) synapses formation and maintenance (Takano et al., 2020; Baldwin et al., 2021; Lee et al., 2021; Wahis et al., 2021b) and can modulate synaptic activity through the release of numerous gliotransmitters (Durkee and Araque, 2019; Goenaga et al., 2023). Since α1-NAR signaling in astrocytes seems to affect some, if not all, of these mechanisms (Wahis and Holt, 2021), it would be interesting to investigate if any of their associated molecular pathways are altered following astrocyte-specific KO of Adra1a, for instance by using a single cell omics approach and look for differential expression of key genes and/or proteins. This might provide interesting targets for future studies aiming at functional validation. However, none of these mechanisms can adequately explain the observation that astrocytic α1A-NAR KD had no effects on the increase in IPSC frequency observed immediately following NA application (Figure 4E). Based on current literature, it appears that this might be best explained by a direct action of NA on GABAergic interneurons, which are another cortical cell type expressing α1A-NAR (Zhang et al., 2014; Zeisel et al., 2015). In fact, there are multiple studies showing that NA application leads to increased interneuron firing and consequently increased inhibition of principal cells in cortex (Bennett et al., 1998; Kawaguchi and Shindou, 1998; Kobayashi et al., 2000) and elsewhere in the CNS (Perez, 2020; Wahis and Holt, 2021), this specifically through α1-NAR signaling and not other NARs (Bergles et al., 1996; Kobayashi et al., 2000; Lei et al., 2007; Dinh et al., 2009). Interestingly, our observation that direct NA effects on IPSC frequency are unaltered following astroglial α1A-NAR KD also indicates that neuronal NAR are not affected by our KD strategy, consistent with our immunostaining data indicating an astrocyte-restricted expression of Adra1a-shRNA (Figure 2). Our observation of a specific role of astrocyte α1A-NAR in modulating the effect of NA on inhibitory circuits is also consistent with the recent literature. Indeed, Reitman and colleagues found that the loss of α1A-NAR signaling in astrocytes (Adra1a cKO) globally increased neuronal activity and network desynchrony. This is consistent with our observation of a lower inhibitory tone and, hence, increased E/I ratio during baseline measures of PSCs in slices from Adra1a-shRNA mice (Figures 4, 5A). Moreover, they also found that astrocyte-specific α1A-NAR cKO alters the fundamental dynamics of NA signaling in cortex, with the impairment of astroglial α1A-NAR signaling impacting the transition of circuits to a more synchronized activity following termination of a period of increased arousal, when phasic release of NA is reduced. This observation correlates well with our observation of a decreased duration of NA effect on IPSC frequency during NA wash out in Adra1a-shRNA mice (Figure 4E). Based on their data, Reitman and colleagues propose that α1A-NAR signaling in astrocytes is important to regulate the magnitude and shape of neuronal responses to NA, probably through the specific modulation of interneuron activity (Reitman et al., 2023), a conclusion consistent with our findings. Indeed, it is not only brain state transitions but also synaptic plasticity that is dependent on the activity of GABAergic interneurons (Calcagnotto, 2016; Tremblay et al., 2016; Zucca et al., 2017; McCormick et al., 2020). When considered alongside other studies, our results are indicative of a dual action of α1-NAR signaling in cortex: it exerts an immediate effect on GABAergic inputs through a direct action on interneurons, while it exerts a slower, modulatory effect on circuit plasticity and the basal parameters of inhibitory inputs through astrocytes. In this context, it is noteworthy that among hippocampal neurons, α1A-NAR seems to be expressed preferentially in SST^+^ interneurons, with NA-induced increases in inhibitory tone in pyramidal CA1 neurons being dependent both on GABA and SST receptors (Hillman et al., 2005, 2009). Future studies should, therefore, delineate the relative participation of astrocytes and (SST) interneurons to α1A-NAR using different Cre-driver lines, to drive gene knock down or gene ablation in these specific cell types.

Our results demonstrate the importance of astrocytic α1-NAR in conveying the neuromodulatory action of NA in a sensory cortex, V1, and provide unique insights into the mechanisms involved. NA signaling is thought to increase the signal to noise ratio of sensory inputs in V1 and other brain regions, notably through its action on inhibitory transmission (Hasselmo et al., 1997; Kobayashi et al., 2000; Polack et al., 2013; Manella et al., 2017). Our study adds a layer of complexity to these mechanisms, by demonstrating an important role for astroglial α1A-NAR in modulating circuit activity in cortex. NA signaling also finely controls the plasticity of cortical synapses, tipping the balance in favor of LTP or LTD depending on the amount of NA released and which NAR are activated, both of which depend on arousal level (Salgado et al., 2016). Our experiments confirm that NA signaling affects plasticity in V1 circuits, and indicate a mechanism in which α1A-NAR signaling, through astrocytes, plays a crucial role in steering the direction of plasticity towards LTP. Indeed, the LTP induction protocol we used, which is known to rely on α1-NAR activation (Pankratov and Lalo, 2015), actually induced LTD in slices with astrocyte-specific α1A-NAR KD (Figure 5). Interestingly, cortical astrocytes have recently been shown to control a developmental switch in plasticity from LTD to LTP in layer IV to layer II-III signaling in the somatosensory cortex (Martínez-Gallego et al., 2022). While the involvement of noradrenergic signaling was not tested in this study (Martínez-Gallego et al., 2022), it clearly adds to the evidence that astrocytes are potent helmsmen of synaptic plasticity direction (Falcón-Moya et al., 2020; Hennes et al., 2020; Wahis et al., 2021b; Koh et al., 2022). Future studies could also explore the molecular mechanisms underlying the role of astrocytes in NA-mediated plasticity in V1, for instance by testing the involvement of potential gliotransmitters, such as ATP/Adenosine, D-Serine, GABA or BDNF (Pankratov and Lalo, 2015; Morioka et al., 2021; Liu et al., 2022; Martínez-Gallego et al., 2022).

Furthermore, we believe it likely that our findings in acute slices extrapolate to the in vivo situation, based on the findings of Monai and colleagues (Monai et al., 2016). In this study, transcranial direct current stimulation (tDCS) was used to potentiate visually evoked potentials in mouse V1. The authors demonstrated that this potentiation is (i) blocked by the α1-NAR antagonist prazosin and (ii) dependent on Ca^2+^ release from intracellular stores through type 2 inositol 1,4,5-trisphosphate receptors (IP_3_R_2_) located in the endoplasmic reticulum of astrocytes. Interestingly, IP_3_R_2_ KO mice fail to show large [Ca^2+^] increases upon α1-NAR activation (Srinivasan et al., 2015). When interpreted alongside our findings, these results strongly suggest that α1A-NAR signaling in astrocytes is instrumental in the potentiation of visual inputs by NA. To confirm this, it would then be interesting to test if the potentiation of visual inputs by tDCS (Monai et al., 2016) is also affected in mice with astroglial α1A-NAR KO.

In conclusion, we found that astrocytic α1A-NAR signaling is a key element of noradrenergic neuromodulation in mice V1. This work reinforces evidence demonstrating that astrocytes are pivotal intermediaries in several other neuromodulatory systems (Papouin et al., 2017; Pacholko et al., 2020; Wahis et al., 2021a), principally the monoaminergic ones (Ma et al., 2016; Kárpáti et al., 2019; Mu et al., 2019; Corkrum et al., 2020; Kohro et al., 2020; Reitman et al., 2023). Our results further strengthen the idea that astrocytes act as signal relays and modulators, in both space and time (Araque et al., 2014), by showing that α1A-NAR signaling in astrocytes is essential both to prolong and potentiate the effect of NA on inhibitory transmission in V1 (Figure 4), with important consequences for plasticity of visual circuits (Figure 5B-E). This work also substantiates the earlier hypothesis that astrocytes might be the principal CNS targets of NA (Salm and McCarthy, 1989, 1992; Stone and Ariano, 1989). In future studies, it will be interesting to determine if α1-NAR signaling through astrocytes has the same consequences for neuromodulation in brain regions other than V1, especially considering that astrocyte heterogeneity, both between and within CNS regions, has emerged as an important factor defining astrocyte functions (Bayraktar et al., 2020; Pestana et al., 2020; Holt, 2023). For instance, earlier work has shown astrocytes in different CNS regions respond to α1-NAR activation, albeit with [Ca^2+^] transients of different kinetics (Batiuk et al., 2020; Pham et al., 2020). However, the consequences of such differences for neuromodulation remain unknown and await clarification. Finally, both the noradrenergic system and astrocytes are involved in the etiology of CNS disorders such as epilepsy (Ghasemi and Mehranfard, 2018; Vezzani et al., 2022) and Alzheimer’s disease (Leanza et al., 2018; Mercan and Heneka, 2022; Perez, 2023). Our work might, therefore, open up new therapeutic avenues, establishing astrocytes as valid alternative targets for therapeutics targeting the NA system.

## ACKNOWLEDGMENTS

We would like to thank Dr. Araks Martirosyan and Mr. Francisco Pestana for advice regarding interpretation of transcriptome-based studies. We would also like to thank Dr. Filip de Vin for help and advice regarding multi electrode array recordings. We also acknowledge the VIB-KU Leuven electrophysiology unit and Dr. Keimpe Wierda for access to electrophysiology setups and advice on experimental design.

## FIGURE LEGENDS

Figure S1. Adra1a-shRNA expression leads to α1A-NAR knockdown

A. Summary of the experimental strategy to test the knockdown efficiency of EGFP-shRNA using a recombinant overexpression system, consisting of an α1A-NAR fused to two V5 epitope tags at the extreme N-terminus of the protein (α1A-V5(x2)-NAR) expressed in HEK293T cells. B. (left) Following fixation and immunocytochemistry, levels of α1A-V5(x2)-NAR were measured cell by cell and averaged per group (right). C. (Left) example western blot image used to quantify the relative amount of α1A- V5(x2)-NAR versus vinculin in cells lysates from each experimental group and (right) bar plot showing levels of α1A-V5(x2)-NAR expression normalized by the vinculin loading control. (B), n = 3 biological replicates per group, 2 coverslips per replicate, p-values displayed above bar plot: Kruskal-Wallis test followed by Dunn’s multiple comparisons. Scale bars: 50 µm. (C) n = 3 biological replicates.

Figure S1 Source data for Western-blot analysis.

Images are taken from the experiments used to quantify α1A-V5(x2)-NAR levels in HEK293T cells lysates (Figure S1). A total of three biological replicates were used. Experimental conditions are indicated at the top of the figure. Red rectangles indicate the bands quantified and used to generate the bar plot in Figure S1C. These bands were chosen based on α1A-NAR predicted weight of 60 kDa (see methods).

Table S1. Descriptive statistics and statistical test parameters

This table provides information on the descriptive statistics of data presented in the figures, as well as information regarding the statistical tests performed in this study. The Shapiro-Wilk or D’Agostino & Pearson normality tests were used with a significance threshold set at 5%. Datasets that were not normally distributed were compared using non-parametric tests.

Table S2. Numbers of astrocytes expressing defined pairs of NARs with the corresponding Spearman correlation test values

Table summarizing the numbers of astrocytes expressing defined pairs of NARs with the corresponding Spearman correlation test values. Based on data from a previously published single cell study (Batiuk et al., 2020) and shown in Figure 1. Some possible pairs of NARs were not found, so were omitted from this table. r: Spearman correlation coefficient.

## SUPPLEMENTARY MATERIALS

Figure S1: EGFP-shRNA expression leads to α1A-NAR knockdown; Figure S1 Source data for Western-blot analysis; Table S1: Descriptive statistics and statistical test parameters; Table S2: Numbers of astrocytes expressing defined pairs of NARs with the corresponding Spearman correlation test values

## AUTHOR CONTRIBUTIONS

JW designed the study with MGH. JW performed the Ca^2+^ imaging and patch-clamp experiments, as well as some of the immunofluorescence experiments. JW also performed and supervised data analysis across experiments (including those performed by other authors). CA performed the in vitro experiments on HEK293T cells. AK, AM, KZ and JV performed stereotaxic injections, as well as some of the IHC experiments (including preliminary analyses). AK and AM performed the MEA experiments, including data extraction from raw signals. HG and SH performed the scRNA-seq data analysis. AK (Northeastern University), AM (Vrije Universiteit Amsterdam), KZ (Northeastern University), HG and SH (Yerevan State University) contributed to this manuscript during the course of their undergraduate studies. MGH conceived and funded the experiments and contributed to data analysis/interpretation. JW wrote the manuscript and designed the figures with input and editing by MGH. All authors approved the final version for submission.

## ETHICS APPROVAL STATEMENT

All experiments were approved by the Ethical Research Committee of the KU Leuven and were in accordance with the European Communities Council Directive of 22 September 2010 (2010/63/EU) and with the relevant Belgian legislation (KB of 29 May 2013). Work presented in this article was covered by Ethische Commissie Dierproeven P226/2018.

## DATA AVAILABILITY STATEMENT

Reagents, analysis scripts and raw data will be made available upon reasonable request to the corresponding authors.

