## Supplementary figures and images for "The astrocyte α1-adrenoreceptor is a key component of the neuromodulatory system in mouse visual cortex"

### Figure S1

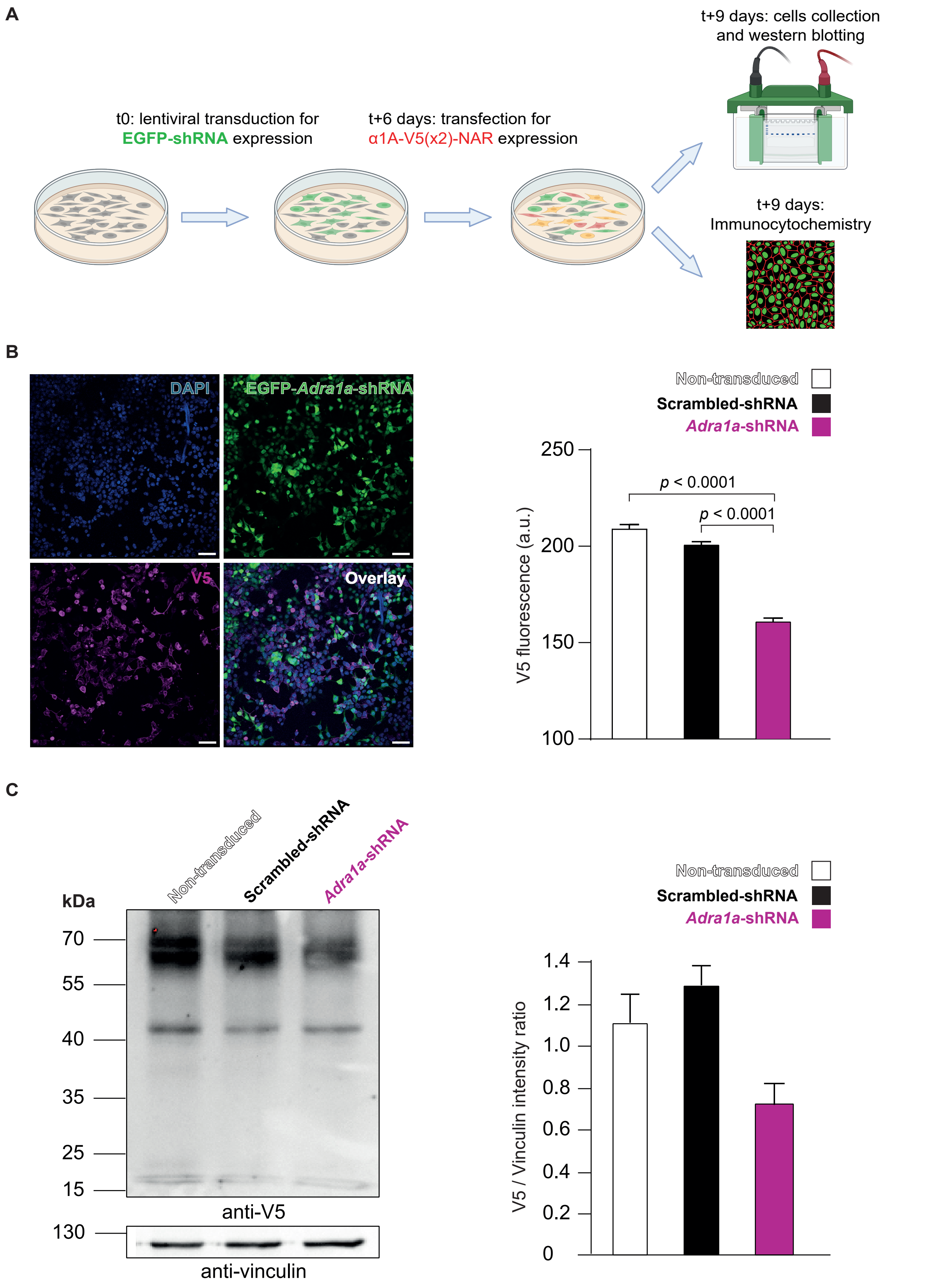


Figure S1 - Wahis et al.,
