## Supplementary material for "The astrocyte α1-adrenoreceptor is a key component of the neuromodulatory system in mouse visual cortex": Figure S1 Source data

### Western-blot image, first set:

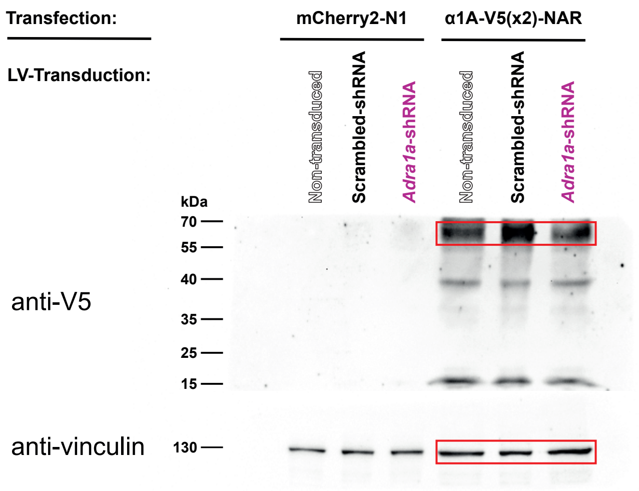


### Western-blot image, second set:

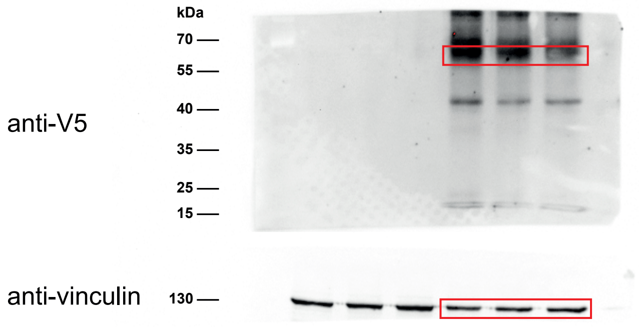


### Western-blot image, third set:

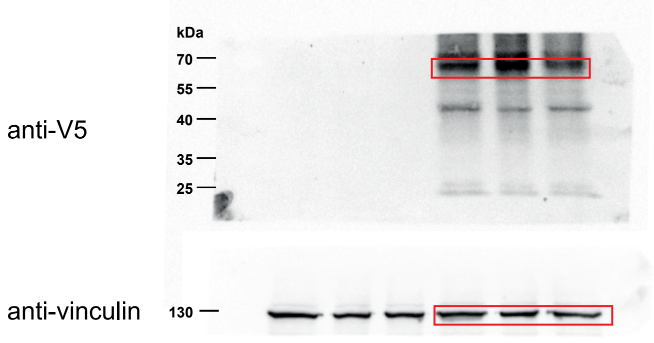


Figure S1 - Source Data - Wahis et al.,
