## Supplementary material for "The astrocyte α1-adrenoreceptor is a key component of the neuromodulatory system in mouse visual cortex": Table S1

| Figure panel data | Group ID & number of replicates | Experimental unit | Variable | Variable unit | N | Mean | SEM | Normality (Shapiro-Wilk Test) ? | Test used | Test statistics | Test *p* -value |
|---|---|---|---|---|---|---|---|---|---|---|---|
| **1** | *Adra1a:* n = 2 littermates (mice) | astrocytes | Normalized expression level | - | 990 | 0.156 | 0.015 | No | | Dunn's test statistic: -12.4430633935899 | 1.220E-34 |
| | *Adra1b:* n = 2 littermates (mice) | | | | 990 | 0.038 | 0.007 | No | | | |
| | *Adra1a:* n = 2 littermates (mice) | | | | 990 | 0.156 | 0.015 | No | | Dunn's test statistic: -17.381655447833 | 9.090E-67 |
| | *Adra1d:* n = 2 littermates (mice) | | | | 990 | 0.002 | 0.001 | No | | | |
| | *Adra1a:* n = 2 littermates (mice) | | | | 990 | 0.156 | 0.015 | No | | Dunn's test statistic: -14.9132384316814 | 2.163E-49 |
| | *Adra2a:* n = 2 littermates (mice) | | | | 990 | 0.020 | 0.005 | No | | | |
| | *Adra1a:* n = 2 littermates (mice) | | | | 990 | 0.156 | 0.015 | No | Kruskal-Wallis test followed by Dunn's multiple comparisons adjusted following the Bonferroni method. Test statistic = 550.23 ; *p* < 0.0001 | Dunn's test statistic: -17.65476569833 | 7.482E-69 |
| | *Adra2b:* n = 2 littermates (mice) | | | | 990 | 0.000 | 0.000 | No | | | |
| | *Adra1a:* n = 2 littermates (mice) | | | | 990 | 0.156 | 0.015 | No | | Dunn's test statistic: -17.65476569833 | 7.482E-69 |
| | *Adra2c:* n = 2 littermates (mice) | | | | 990 | 0.000 | 0.000 | No | | | |
| | *Adra1a:* n = 2 littermates (mice) | | | | 990 | 0.156 | 0.015 | No | | Dunn's test statistic: -10.2775196309538 | 7.119E-24 |
| | *Adrb1:* n = 2 littermates (mice) | | | | 990 | 0.044 | 0.007 | No | | | |
| | *Adra1a:* n = 2 littermates (mice) | | | | 990 | 0.156 | 0.015 | No | | Dunn's test statistic: -17.2493103224934 | 9.059E-66 |
| | *Adrb2:* n = 2 littermates (mice) | | | | 990 | 0.001 | 0.001 | No | | | |
| | *Adra1a:* n = 2 littermates (mice) | | | | 990 | 0.156 | 0.015 | No | | Dunn's test statistic: -17.65476569833 | 7.482E-69 |
| | *Adrb3:* n = 2 littermates (mice) | | | | 990 | 0.000 | 0.000 | No | | | |
| **2B** | *S100beta positive cells (n = 4 mice)* | average % per section | Proportion of cells | | 2399 | | | | | | |
| | *S100beta and EGFP positive cells (n = 4 mice)* | | | ratio | 859 | 0.358 | 0.052 | | | | |
| | *NeuN positive cells (n = 4 mice)* | | | | 10446 | | | | | | |
| | *NeuN and EGFP positive cells (n = 4 mice)* | | | ratio | 22 | 0.002 | 0.001 | | | | |
| **3D** | Scrambled-shRNA: n = 8 mice | astrocytes | ΔF/F0 Amplitude | % | 38 | 203.964 | 13.967 | No | Two-sided Wilcoxon rank sum test | zval = -0.762350880233423 | 4.459E-01 |
| | *Adra1a-* shRNA: n = 7 mice | | ΔF/F0 Amplitude | % | 70 | 181.412 | 6.983 | No | | ranksum = 3696 | |
| | Scrambled-shRNA: n = 8 mice | | Time of Peak ΔF/F0 | s | 38 | 149.333 | 10.322 | No | Two-sided Wilcoxon rank sum test | zval =3.36145257191589 | 7.753E-04 |
| | *Adra1a-* shRNA: n = 7 mice | | Time of Peak ΔF/F0 | s | 70 | 166.955 | 6.002 | No | | ranksum = 4338 | |
| | Scrambled-shRNA: n = 8 mice | | ΔF/F0 AUC | %.s | 38 | 449.373 | 24.675 | No | Two-sided Wilcoxon rank sum test | zval = -0.395650456830004 | 6.924E-01 |
| | *Adra1a-* shRNA: n = 7 mice | | ΔF/F0 AUC | %.s | 70 | 414.343 | 10.594 | No | | ranksum = 3753 | |
| **3E** | Scrambled-shRNA: n = 8 mice | astrocytes | ΔF/F0 Amplitude | % | 49 | 173.361 | 8.352 | No | Two-sided Wilcoxon rank sum test | zval = -2.14951600407325 | 3.159E-02 |
| | *Adra1a-* shRNA: n = 7 mice | | ΔF/F0 Amplitude | % | 32 | 146.767 | 7.853 | No | | ranksum = 1089 | |
| | Scrambled-shRNA: n = 8 mice | | Time of Peak ΔF/F0 | s | 49 | 146.351 | 10.114 | No | Two-sided Wilcoxon rank sum test | zval = 4.16861193598925 | 3.065E-05 |
| | *Adra1a-* shRNA: n = 7 mice | | Time of Peak ΔF/F0 | s | 32 | 202.187 | 10.315 | Yes | | ranksum = 1744 | |
| | Scrambled-shRNA: n = 8 mice | | ΔF/F0 AUC | %.s | 49 | 402.293 | 13.844 | No | Two-sided Wilcoxon rank sum test | zval = -2.13019451190181 | 3.316E-02 |
| | *Adra1a-* shRNA: n = 7 mice | | ΔF/F0 AUC | %.s | 32 | 356.616 | 10.475 | No | | ranksum = 1091 | |
| **4E** | Scrambled-shRNA: n = 6 mice | neurons | IPSC frequency baseline | Hz | 20 | 1.257 | 0.209 | No | Two-sided Wilcoxon rank sum test | zval = 2.5988 | 9.400E-03 |
| | *Adra1a-* shRNA: n = 5 mice | | IPSC frequency baseline | Hz | 13 | 0.674 | 0.192 | No | | ranksum = 411 | |
| | Scrambled-shRNA: n = 6 mice | | IPSC frequency NA Wash in | Hz | 20 | 1.900 | 0.313 | No | Two-sided Wilcoxon rank sum test | zval = 1.5661 | 1.173E-01 |
| | *Adra1a-* shRNA: n = 5 mice | | IPSC frequency NA Wash in | Hz | 13 | 1.432 | 0.413 | No | | ranksum = 383 | |
| | Scrambled-shRNA: n = 6 mice | | IPSC frequency NA Wash out | Hz | 20 | 1.588 | 0.363 | No | Two-sided Wilcoxon rank sum test | zval = 2.1382 | 3.250E-02 |
| | *Adra1a-* shRNA: n = 5 mice | | IPSC frequency NA Wash out | Hz | 13 | 0.772 | 0.206 | No | | ranksum = 398.5000 | |
| | Scrambled-shRNA: n = 6 mice | | IPSC frequency baseline | Hz | 20 | 1.257 | 0.209 | No | Two-sided Wilcoxon signed rank test | zval = -2.6326 | 8.500E-03 |
| | Scrambled-shRNA: n = 6 mice | | IPSC frequency NA Wash in | Hz | 20 | 1.900 | 0.313 | No | | signed rank = 34.5000 | |
| | Scrambled-shRNA: n = 6 mice | | IPSC frequency NA Wash out | Hz | 20 | 1.588 | 0.363 | No | Two-sided Wilcoxon signed rank test | zval = -1.4085 | 1.590E-01 |
| | Scrambled-shRNA: n = 6 mice | | IPSC frequency NA Wash in | Hz | 20 | 1.900 | 0.313 | No | | signed rank = 60 | |
| | *Adra1a-* shRNA: n = 5 mice | | IPSC frequency baseline | Hz | 13 | 0.674 | 0.192 | No | Two-sided Wilcoxon signed rank test | | 4.883E-04 |
| | *Adra1a-* shRNA: n = 5 mice | | IPSC frequency NA Wash in | Hz | 13 | 1.432 | 0.413 | No | | signed rank = 1 | |
| | *Adra1a-* shRNA: n = 5 mice | | IPSC frequency NA Wash out | Hz | 13 | 0.772 | 0.206 | No | Two-sided Wilcoxon signed rank test | | 2.700E-03 |
| | *Adra1a-* shRNA: n = 5 mice | | IPSC frequency NA Wash in | Hz | 13 | 1.432 | 0.413 | No | | signed rank = 5.5000 | |
| **4F** | Scrambled-shRNA: n = 6 mice | neurons | IPSC amplitude baseline | pA | 20 | 27.158 | 1.606 | Yes | Two-sided Wilcoxon rank sum test | zval = -0.8658 | 3.866E-01 |
| | *Adra1a-* shRNA: n = 5 mice | | IPSC amplitude baseline | pA | 13 | 36.248 | 6.985 | No | | ranksum = 316 | |
| | Scrambled-shRNA: n = 6 mice | | IPSC amplitude NA Wash in | pA | 20 | 32.630 | 3.461 | No | Two-sided Wilcoxon rank sum test | zval = 0 | 1.000E+00 |
| | *Adra1a-* shRNA: n = 5 mice | | IPSC amplitude NA Wash in | pA | 13 | 33.958 | 4.812 | Yes | | ranksum = 340 | |
| | Scrambled-shRNA: n = 6 mice | | IPSC amplitude NA Wash out | pA | 20 | 29.168 | 3.505 | No | Two-sided Wilcoxon rank sum test | zval = -0.0553 | 9.559E-01 |
| | *Adra1a-* shRNA: n = 5 mice | | IPSC amplitude NA Wash out | pA | 13 | 27.898 | 2.561 | Yes | | ranksum = 338 | |
| | Scrambled-shRNA: n = 6 mice | | IPSC amplitude baseline | pA | 20 | 27.158 | 1.606 | Yes | Two-sided Wilcoxon signed rank test | zval = -2.0160 | 4.380E-02 |
| | Scrambled-shRNA: n = 6 mice | | IPSC amplitude NA Wash in | pA | 20 | 32.630 | 3.461 | No | | signed rank = 51 | |
| | Scrambled-shRNA: n = 6 mice | | IPSC amplitude NA Wash out | pA | 20 | 29.168 | 3.505 | No | Two-sided Wilcoxon signed rank test | zval = -1.6800 | 9.300E-02 |
| | Scrambled-shRNA: n = 6 mice | | IPSC amplitude NA Wash in | pA | 20 | 32.630 | 3.461 | No | | signed rank = 60 | |
| | *Adra1a-* shRNA: n = 5 mice | | IPSC amplitude baseline | pA | 13 | 36.248 | 6.985 | No | Two-sided Wilcoxon signed rank test | | 8.926E-01 |
| | *Adra1a-* shRNA: n = 5 mice | | IPSC amplitude NA Wash in | pA | 13 | 33.958 | 4.812 | Yes | | signed rank = 43 | |
| | *Adra1a-* shRNA: n = 5 mice | | IPSC amplitude NA Wash out | pA | 13 | 27.898 | 2.561 | Yes | Paired *t-* test | Confidence interval = [-13.7262, 1.6056] | 1.106E-01 |
| | *Adra1a-* shRNA: n = 5 mice | | IPSC amplitude NA Wash in | pA | 13 | 33.958 | 4.812 | Yes | | *t* stat = -1.7225 | |
| | | | | | | | | | | dF = 12 | |
| | | | | | | | | | | SD = 12.6857 | |
| **4G** | Scrambled-shRNA: n = 5 mice | neurons | EPSC frequency baseline | Hz | 8 | 1.483 | 0.358 | No | Two-sided Wilcoxon rank sum test | | 2.667E-01 |
| | *Adra1a-* shRNA: n = 4 mice | | EPSC frequency baseline | Hz | 8 | 1.065 | 0.227 | No | | ranksum = 79 | |
| | Scrambled-shRNA: n = 5 mice | | EPSC frequency NA Wash in | Hz | 8 | 2.271 | 0.665 | No | Two-sided Wilcoxon rank sum test | | 6.500E-02 |
| | *Adra1a-* shRNA: n = 4 mice | | EPSC frequency NA Wash in | Hz | 8 | 1.513 | 0.286 | No | | ranksum = 86 | |
| | Scrambled-shRNA: n = 5 mice | | EPSC frequency NA Wash out | Hz | 8 | 1.360 | 0.132 | Yes | Two-sided Wilcoxon rank sum test | | 1.049E-01 |
| | *Adra1a-* shRNA: n = 4 mice | | EPSC frequency NA Wash out | Hz | 8 | 1.123 | 0.275 | No | | ranksum = 84 | |
| | Scrambled-shRNA: n = 5 mice | | EPSC frequency baseline | Hz | 8 | 1.483 | 0.358 | No | Two-sided Wilcoxon signed rank test | | 1.484E-01 |

| | | | | | | | | | | | |
|---|---|---|---|---|---|---|---|---|---|---|---|
| | Scrambled-shRNA: n = 5 mice | | EPSC frequency NA Wash in | Hz | 8 | 2.271 | 0.665 | No | | signed rank = 7 | |
| | Scrambled-shRNA: n = 5 mice | | EPSC frequency NA Wash out | Hz | 8 | 1.360 | 0.132 | Yes | Two-sided Wilcoxon signed rank test | | 1.172E-01 |
| | Scrambled-shRNA: n = 5 mice | | EPSC frequency NA Wash in | Hz | 8 | 2.271 | 0.665 | No | | signed rank = 6.5 | |
| | *Adra1a*- shRNA: n = 4 mice | | EPSC frequency baseline | Hz | 8 | 1.065 | 0.227 | No | Two-sided Wilcoxon signed rank test | | 1.094E-01 |
| | *Adra1a*- shRNA: n = 4 mice | | EPSC frequency NA Wash in | Hz | 8 | 1.513 | 0.286 | No | | signed rank = 6 | |
| | *Adra1a*- shRNA: n = 4 mice | | EPSC frequency NA Wash out | Hz | 8 | 1.123 | 0.275 | No | Two-sided Wilcoxon signed rank test | | 7.810E-02 |
| | *Adra1a*- shRNA: n = 4 mice | | EPSC frequency NA Wash in | Hz | 8 | 1.513 | 0.286 | No | | signed rank = 5 | |
| **4H** | Scrambled-shRNA: n = 5 mice | neurons | EPSC amplitude baseline | pA | 8 | 19.046 | 2.662 | Yes | Independent sample *t*- test | Confidence interval = [-6.9969, 5.5942] | 8.146E-01 |
| | *Adra1a*- shRNA: n = 4 mice | | EPSC amplitude baseline | pA | 8 | 18.345 | 1.236 | Yes | | *t* stat = -0.2389 | |
| | | | | | | | | | | dF = 14 | |
| | | | | | | | | | | SD = 5.8706 | |
| | Scrambled-shRNA: n = 5 mice | | EPSC amplitude NA Wash in | pA | 8 | 21.277 | 2.354 | No | Two-sided Wilcoxon rank sum test | | 9.591E-01 |
| | *Adra1a*- shRNA: n = 4 mice | | EPSC amplitude NA Wash in | pA | 8 | 23.810 | 3.613 | Yes | | ranksum = 67 | |
| | Scrambled-shRNA: n = 5 mice | | EPSC amplitude NA Wash out | pA | 8 | 20.337 | 3.737 | No | Two-sided Wilcoxon rank sum test | | 8.785E-01 |
| | *Adra1a*- shRNA: n = 4 mice | | EPSC amplitude NA Wash out | pA | 8 | 18.815 | 2.043 | Yes | | ranksum = 66 | |
| | Scrambled-shRNA: n = 5 mice | | EPSC amplitude baseline | pA | 8 | 19.046 | 2.662 | Yes | Two-sided Wilcoxon signed rank test | | 3.125E-01 |
| | Scrambled-shRNA: n = 5 mice | | EPSC amplitude NA Wash in | pA | 8 | 21.277 | 2.354 | No | | signed rank = 10 | |
| | Scrambled-shRNA: n = 5 mice | | EPSC amplitude NA Wash out | pA | 8 | 20.337 | 3.737 | No | Two-sided Wilcoxon signed rank test | | 3.828E-01 |
| | Scrambled-shRNA: n = 5 mice | | EPSC amplitude NA Wash in | pA | 8 | 21.277 | 2.354 | No | | signed rank = 11 | |
| | *Adra1a*- shRNA: n = 4 mice | | EPSC amplitude baseline | pA | 8 | 18.345 | 1.236 | Yes | Paired *t*- test | Confidence interval = [-13.6529, 2.7216] | 1.584E-01 |
| | *Adra1a*- shRNA: n = 4 mice | | EPSC amplitude NA Wash in | pA | 8 | 23.810 | 3.613 | Yes | | *t* stat = -1.5786 | |
| | | | | | | | | | | dF = 7 | |
| | | | | | | | | | | SD = 9.7932 | |
| | *Adra1a*- shRNA: n = 4 mice | | EPSC amplitude NA Wash out | pA | 8 | 18.815 | 2.043 | Yes | Paired *t*- test | Confidence interval = [-4.0780, 14.0689] | 2.342E-01 |
| | *Adra1a*- shRNA: n = 4 mice | | EPSC amplitude NA Wash in | pA | 8 | 23.810 | 3.613 | Yes | | *t* stat = 1.3019 | |
| | | | | | | | | | | dF = 7 | |
| | | | | | | | | | | SD = 10.8531 | |
| **5E** | Scrambled-shRNA: n = 3 mice | Slices | fEPSP amplitude | Δ% baseline | 6 | 28.410 | 2.335 | Yes | Independent sample *t*- test | Confidence interval = [26.6610, 44.5397] | 8.469E-06 |
| | *Adra1a*- shRNA: n = 3 mice | | fEPSP amplitude | Δ% baseline | 5 | -7.192 | 3.315 | Yes | | *t* stat = 9.0089 | |
| | | | | | | | | | | dF = 9 | |
| | | | | | | | | | | SD = 6.5260 | |
| **S1B** | Non-transduced: n = 3 biological replicates and 2 coverslips per replicate | HEK293T cells | Absolute fluorescence of V5 channel | a.u. | 6911 | 209.900 | 2.423 | No (D'Agostino & Pearson test) | Kruskal-Wallis test followed by Dunn's multiple comparisons. Test statistic = 828.1 ; *p* < 0.0001 | z = 1.644 | 0.3005 |
| | Scrambled-shRNA: n = 3 biological replicates and 2 coverslips per replicate | HEK293T cells | Absolute fluorescence of V5 channel | a.u. | 8873 | 201.700 | 1.829 | No (D'Agostino & Pearson test) | | | |
| | Non-transduced: n = 3 biological replicates and 2 coverslips per replicate | HEK293T cells | Absolute fluorescence of V5 channel | a.u. | 6911 | 209.900 | 2.423 | No (D'Agostino & Pearson test) | | z = 23.40 | < 0.0001 |
| | *Adra1a* -shRNA: n = 3 biological replicates and 2 coverslips per replicate | HEK293T cells | Absolute fluorescence of V5 channel | a.u. | 7153 | 161.800 | 1.965 | No (D'Agostino & Pearson test) | | | |
| | Scrambled-shRNA: n = 3 biological replicates and 2 coverslips per replicate | HEK293T cells | Absolute fluorescence of V5 channel | a.u. | 8873 | 201.700 | 1.829 | No (D'Agostino & Pearson test) | | z = 26.50 | < 0.0001 |
| | *Adra1a* -shRNA: n = 3 biological replicates and 2 coverslips per replicate | HEK293T cells | Absolute fluorescence of V5 channel | a.u. | 7153 | 161.800 | 1.965 | No (D'Agostino & Pearson test) | | | |
| **S1C** | Non-transduced: n = 3 biological replicates | HEK293T whole cell lysate | Ratio of anti-V5 and anti-Vinculin band intensities | ratio | 3 | 1.111 | 0.139 | | | | |
| | Scrambled-shRNA: n = 3 biological replicates | HEK293T whole cell lysate | Ratio of anti-V5 and anti-Vinculin band intensities | ratio | 3 | 1.294 | 0.090 | | | | |
| | *Adra1a* -shRNA: n = 3 biological replicates | HEK293T whole cell lysate | Ratio of anti-V5 and anti-Vinculin band intensities | ratio | 3 | 0.723 | 0.104 | | | | |
