## Supplementary material for "The astrocyte α1-adrenoreceptor is a key component of the neuromodulatory system in mouse visual cortex": Table S2

| Co-expressed NAR pairs | Number of astrocytes co-expressing defined NAR pair | r | *p* -value |
|---|---|---|---|
| *Adra1a - Adra1b* | 6 | 0.012 | 0.7 |
| *Adra1a - Adra1d* | 1 | 0.047 | 0.14 |
| *Adra1a - Adrb1* | 9 | 0.03 | 0.34 |
| *Adra1b - Adra2a* | 1 | 0.0093 | 0.77 |
| *Adra1b - Adrb1* | 4 | 0.046 | 0.15 |
| *Adra2a - Adrb1* | 1 | -0.0036 | 0.91 |

Wahis et al., Table S2
